# Arabidopsis–Swede Midge as an experimentally tractable insect gall model to dissect plant reprogramming

**DOI:** 10.64898/2026.10.11.758373

**Authors:** Zeynep Begüm Şen, Zsuzsanna Mérai, Marina Silvestre Vaňó, Byron Rusnak, Carina Lopez, Paige Guevarra, Liya Zaygerman, Boyd A Mori, Yan Ma

## Abstract

Insect-induced galls are striking examples of host reprogramming, yet their causal mechanisms remain poorly understood because few insect gall systems combine experimental tractability with precise control of insect and host developmental state. To address this gap, we establish *Arabidopsis thaliana* and the gall-inducing Swede Midge (*Contarinia nasturtii*) as an experimental system for insect gall research.

We discovered that the Swede Midge, an invasive pest of Brassica crops, completes its full life cycle on Arabidopsis, and induces characteristic inflorescence galls resembling those on crops.

A larval phenotyping pipeline monitors how many larvae are present and how far they have developed, and a larval droplet assay delivers defined numbers of stage-selected larvae directly to host, fixing the insect input and establishing a precise time zero for the interaction. We found that early-stage, but not late-stage, larvae induce galls.

A phyllotactic dissection and positional annotation workflow enables comparisons of galled and non-galled floral tissues. We found that galling enlarged floral buds, floral primordia and the inflorescence meristem while delaying floral developmental progression, with accompanying changes in cell size, shape and tissue organisation.

These resources combine Arabidopsis genetics with controlled insect manipulation, opening the way to mechanistic dissection of how insects reprogram plant pathways.

## Introduction

Galling insects hijack plant cellular machinery to induce specialised structures with protective and nutritional benefits at the expense of plant health. Although made entirely from plant tissue, these ectopic structures can differ markedly from normal host organs. They are often highly organised and morphologically diverse, with forms unique to particular insect-host associations (Stone & Schönrogge, 2003; Harris & Pitzschke, 2020), unlike the relatively unpatterned crown galls induced by *Agrobacterium tumefaciens* (Gohlke & Deeken, 2014). Such specificity has led galls to be regarded as extended phenotypes of the inducing insect (Raman, 2011; Pan *et al*., 2015; Gatjens-Boniche, 2019; Favery *et al*., 2020).

Gall development is closely coordinated with insect development, and many gall-forming insects are obligate parasites that depend on successful gall induction to complete their life cycle (Oliveira *et al*., 2016; Ma *et al*., 2025). At the cellular level, these insects can redirect existing host cells towards new fates and generate specialised tissues tailored to the insect’s nutritional and protective needs (Ma *et al*., 2025). Such extensive reprogramming offers a powerful system for understanding how plant developmental programmes can be reshaped by external biological signals.

Many gall-inducing insects are major agricultural pests, including Phylloxera on grapevine (Granett *et al*., 2001) and Hessian Fly on wheat (Stuart *et al*., 2012). Despite their importance and extensive descriptive, ecological and molecular studies, we still know little about the causal mechanisms that determine insect gall formation. Advances in insect genomics and tissue- or stage-specific transcriptomics have recently enabled identification of candidate insect effectors (Aggarwal *et al*., 2014; Aljbory *et al*., 2020; Korgaonkar *et al*., 2021; Hirano *et al*., 2025). Comparable omics studies in the host have identified many gall-associated pathways, particularly those involving phytohormone signalling, defence and metabolism (Takeda *et al*., 2019; Yang *et al*., 2023; Shih *et al*., 2025; Schultz & Stone, 2026)

Association, however, does not establish mechanism: most insect gall systems lack the genetic tools needed to perturb candidate insect effectors or host targets and test how they causally influence gall formation. Gall-inducing insects are often difficult to rear and manipulate because of complex life cycles and host requirements, while their host plants are frequently non-model species with limited genetic resources and transformation tools. The scarcity of experimentally tractable models therefore remains a major barrier to causal analysis of insect gall formation.

A related challenge is precise developmental staging of both insect and host. Galling is a coupled developmental process involving two changing organisms: its outcome depends simultaneously on insect-derived inputs and the developmental state of the host tissue. In many insect gall systems, particularly those relying on field-collected material, the onset of gall induction and the developmental history of both partners are difficult to determine. Consequently, samples collected at the same nominal time may represent different biological stages, reducing reproducibility and the interpretability of molecular and phenotypic data. Mechanistic analysis therefore requires systems in which insect and host developmental variation can be controlled and accounted for.

We establish a model combining *Arabidopsis thaliana* with the gall-inducing Swede Midge (*Contarinia nasturtii,* **SM**). Arabidopsis provides a genetically tractable host with well-characterised development, extensive mutant libraries and a growing toolkit for gene editing and cell-type-specific reporters (Lee *et al*., 2025; Strader *et al*., 2025). Swede Midge is a Brassicaceae specialist belonging to the Cecidomyiidae (Diptera), a major family of gall-inducing insects (Tokuda, 2026). Native to Europe, Swede Midge has a broad host range including cultivated and wild Brassicaceae and has become an economically important invasive pest in North America (Hallett & Heal, 2001; Chen *et al*., 2009, 2011; Lopez *et al*., 2026). Swede Midge has a short life cycle and established laboratory-rearing methods (Readshaw, 1966; Boddum *et al*., 2018; Mori *et al*., 2024), together with a compact genome and published developmental and organ-specific transcriptomes (Mori *et al*., 2021, 2023, 2024). SM therefore offers a combination of ecological relevance, economic importance and experimental accessibility that makes it particularly well suited as a model gall-inducing insect that complements Arabidopsis.

Here, we establish the Arabidopsis-Swede Midge system as an experimentally tractable resource for insect gall research. We developed complementary approaches to quantify gall formation while controlling and accounting for developmental variation in both partners. Using this system, we identified the first (L1) and second (L2), but not the third (L3), larval instars as competent insect stages to induce galls. We determined that host genetic differences affect both insect preference and gall development. We also found that galling enlarged floral buds, primordia and the inflorescence meristem, delayed floral progression and altered cell morphology and tissue organisation. Our framework now provides the experimental precision needed to compare galling outcomes across genotypes and treatments.

## Results

### 1. Swede Midge completes its life cycle and induces inflorescence galls on *Arabidopsis thaliana*

Swede Midge (SM) is a gall-inducing Brassicaceae specialist and major pest of crops including oilseed rape (*Brassica napus*), broccoli and cauliflower (*Brassica oleracea* var. *italica* and *botrytis*, respectively) (Hallett, 2007). Although its host range is broad within the Brassicaceae (Chen *et al*., 2009; Lopez *et al*., 2026), successful oviposition and larval development on Arabidopsis have only recently been reported (Lopez *et al*., 2026), while its complete life cycle and gall-inducing capacity on this host have not yet been characterised. We therefore tested whether lab-reared SM could infest *Arabidopsis thaliana* (*At*), induce galls and complete its life cycle under controlled conditions.

We established laboratory SM colonies maintained on *Brassica napus* (*Bn*, oilseed rape) and compared SM development on *At*, Rapid Cycling *Brassica rapa* (*Br*) and *Bn*. We demonstrated in two independent laboratories that SM can complete its life cycle on Arabidopsis in 20-30 days for non-diapausing individuals. Developmental timing closely matched that on *Br* and *Bn* under comparable conditions (**Fig.1A**).

**Fig. 1.**
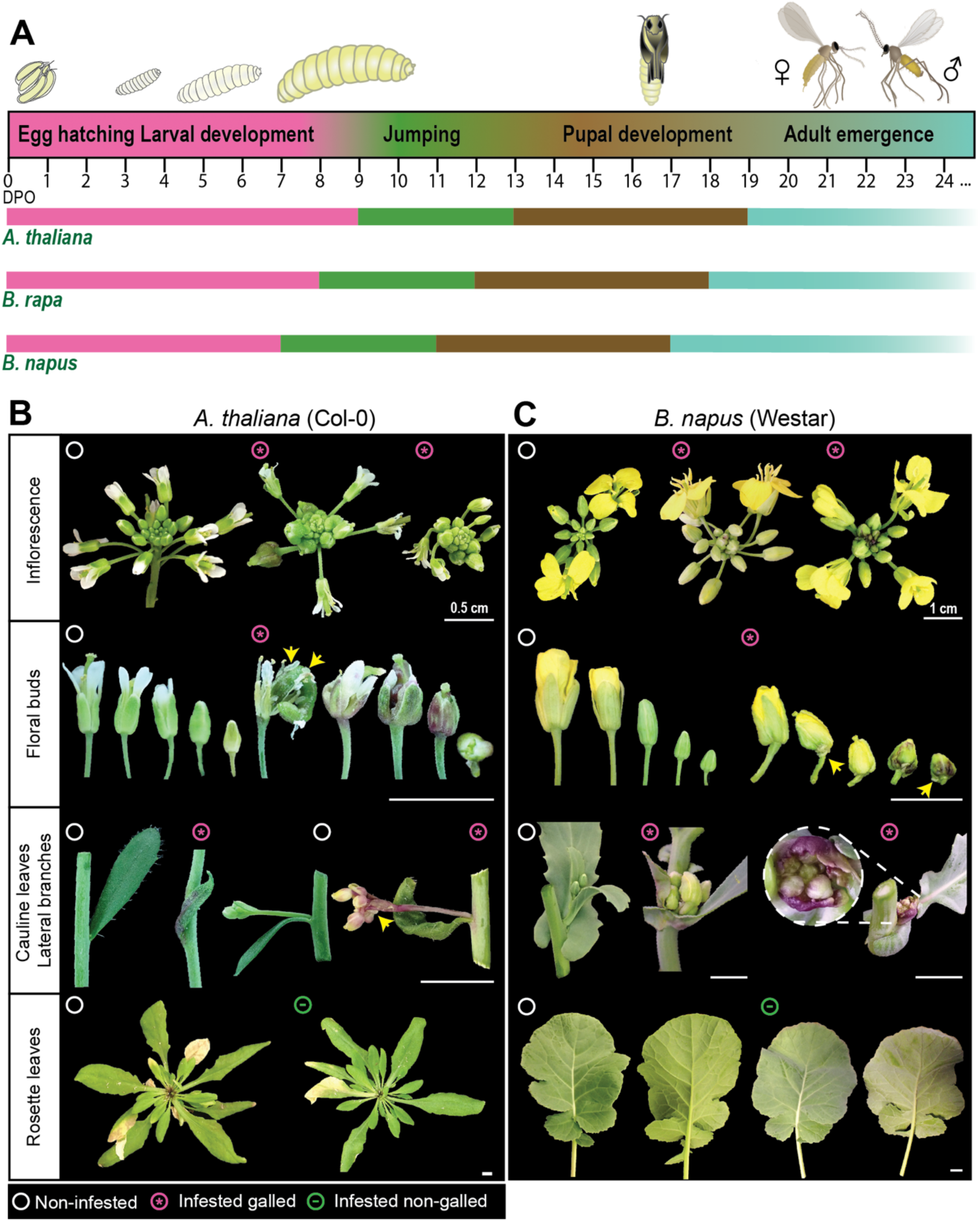
Swede Midge life cycle on Brassicaceae host species and associated gall symptoms. **(A)** Schematics of life cycles on *A. thaliana*, Rapid Cycling *B. rapa* and *B. napus* under lab conditions. Developmental phases include egg hatching and larval development (L1–L3) on the host plant, mature 3^rd^ instar larvae (L3) jumping from the plant, pupal development in soil, adult emergence and subsequent oviposition. Numbers indicate days post-oviposition (DPO). Boundaries for L3 jumping and adult emergence mark the first days of consistent observation. Insect depictions were rendered from real photos and microscopy images (see Fig. S1). Representative gall symptoms at 7–8 DPO are shown for **(B)** *A. thaliana* (Col-0) and **(C)** *B. napus* (Westar) following whole-plant exposure to Swede Midge adults. Non-infested controls, infested galled tissues, and infested non-galled tissues are shown with the indicated symbols. Arrows mark visible larvae on plant tissues. Scale bars = 0.5 cm **(B)** and 1 cm **(C)**.

Adult SM are short-lived, with males typically surviving for 1-3 days and females sometimes persisting for several days longer. Females oviposit on young host tissues, and egg clusters could be observed on young inflorescence tissues after removal of the older outer buds (**Fig. S3C**). The resulting larvae feed within growing galls and develop through three instars (L1-L3), before mature third instar (L3) larvae jump from the plant to pupate in soil and subsequently emerge as the next generation of adults (**Fig. S1**). Mature L3 larvae began leaving plants at ∼9 days post oviposition (DPO) on *At*, compared with ∼8 DPO on *Br* and ∼7 DPO on *Bn* (**Fig. 1A**). Peak adult emergence occurred at 21, 20 and 19 DPO on *At*, *Br* and *Bn*, respectively (**Fig. S2A**).

Equal infestation with 50 male and 50 female adults per cage produced significantly fewer next-generation adults on *At* (75 ± 14 per cage, mean ± SE) than on *Br* (368 ± 36) or *Bn* (551 ± 132), whereas *Br* and *Bn* did not differ significantly (**Fig. S2B**). *At* therefore supported completion of the SM life cycle but yielded fewer next-generation adults than the two other Brassicaceae hosts tested. Representative images of *At*, *Br* and *Bn* at early flowering stages used for infestation also illustrate marked differences in overall plant size and architecture (**Fig. S2C**).

SM infestation produced conspicuous gall phenotypes on *At* that resembled those observed on *Bn* and *Br* (**Fig. 1B, C, S3A, B**). Galled inflorescences were compacted and contained enlarged, deformed floral buds with suppressed flower opening, together with thickening and distortion of floral organs and supporting tissues. Twisting of stems or pedicels and purple pigmentation were also frequently observed (**Fig. 1B, C, S3A, B**). Larvae were observed on young floral buds and within closed buds (**Fig. 1B, C, S8A**).

SM gall formation showed a distinct tissue distribution on *At*. Following whole-plant exposure to SM adults, galls formed predominantly on the main and lateral inflorescences and occasionally on cauline leaves, whereas rosette leaves remained non-galled (**Fig. 1B**). A similar distribution was observed on *Bn* and *Br*, where galling was concentrated around inflorescences and developing floral tissues (**Fig. 1C, S3**). These results suggest that actively developing reproductive tissues are the principal sites of SM gall formation under our lab conditions.

We establish Arabidopsis as a relevant experimental host for SM, supporting its complete life cycle and characteristic inflorescence gall phenotypes broadly comparable to those on agriculturally important Brassicaceae hosts.

### 2. Controlled infestation assays reveal quantitative differences in gall induction and larval performance between Arabidopsis ecotypes

To identify Arabidopsis developmental stages most suitable for SM infestation, we first developed a whole-cage assay in which newly emerged SM adults were released into cages and allowed access to entire plants (**Fig. 2A**). Oviposition depended strongly on *At* developmental stage: with 8 males and 8 females (8M+8F), no eggs were detected on vegetative-stage plants (1.14), but eggs were present at budding (5.10) and early flowering (6.10) stages (Boyes *et al*., 2001). Increasing adult input to 20M+20F generally increased oviposition, most clearly at the early flowering stage (**Fig. S4A, B**). These results identified reproductive-stage plants as suitable for oviposition, and we therefore used *At* plants with actively developing flowers for subsequent whole-cage infestations. Under these conditions, exposure to 50M+50F produced characteristic inflorescence galls in both Col-0 and Ler-0 by 8 DPO (**Fig. 2B**).

**Fig. 2.**
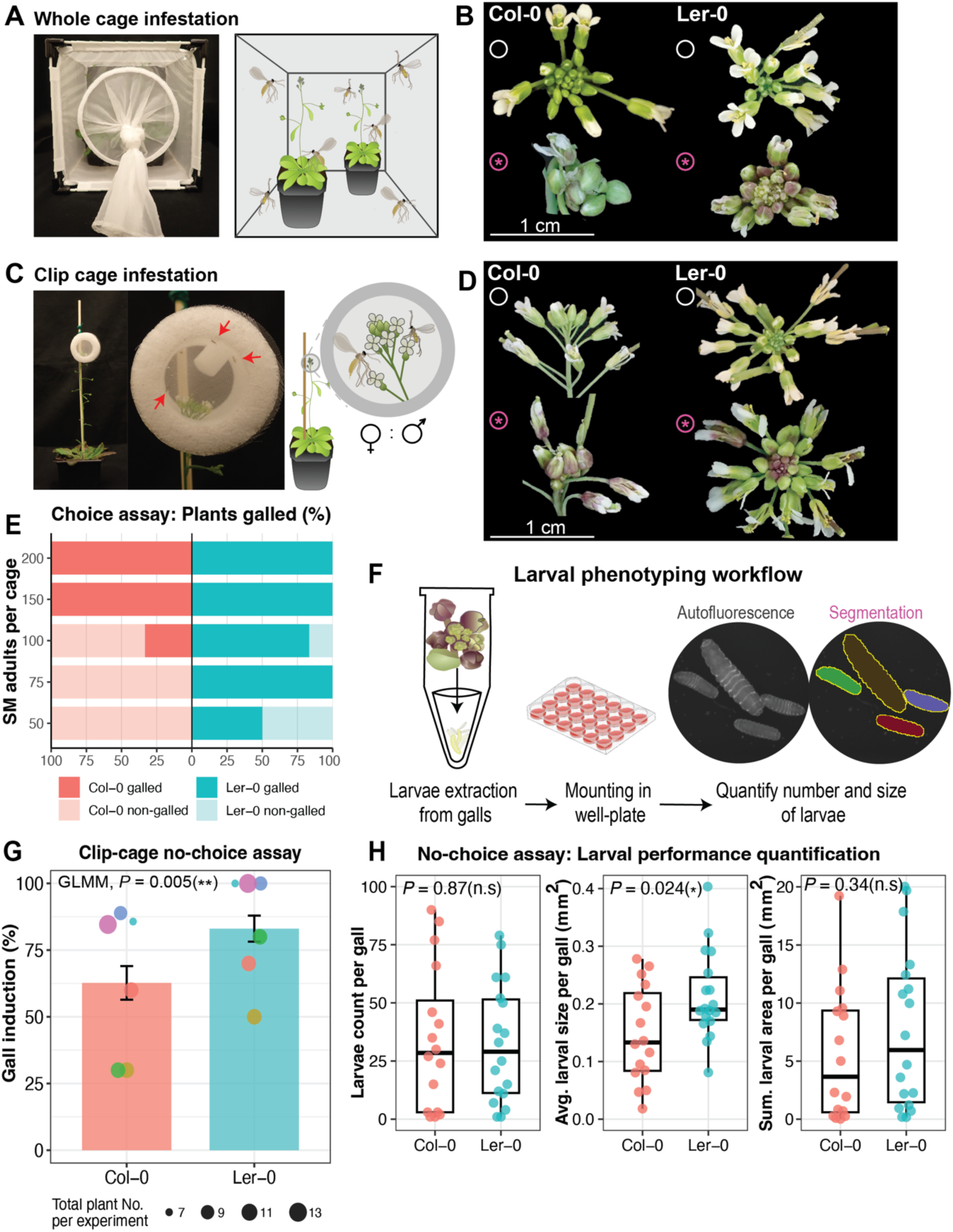
Arabidopsis infestation protocols and susceptibility assays. **(A)** Whole-cage infestation. For no-choice assays, two plants of the same genotype were exposed together within a cage; for choice assays, one plant from each genotype was co-exposed within a cage. **(B)** Representative Col-0 and Ler-0 inflorescences following whole-cage no-choice infestation with 50M+50F adults, assessed at 8 DPO. White circles indicate controls; pink stars indicate infested and galled samples. Scale bars = 1 cm. **(C)** Clip-cage infestation. 5M+5F adults were confined for 24 h to a defined region of the plant, including individual main or lateral inflorescences. **(D)** Representative Col-0 and Ler-0 inflorescences following clip-cage infestation of the main inflorescence with 5M+5F adults per plant, assessed at 7 DPO. White circles indicate controls; pink stars indicate infested and galled samples. Scale bars = 1 cm. **(E)** Whole-cage choice assay comparing gall induction percentages in Col-0 and Ler-0 at 7 DPO. Stacked bars show the percentage of plants that were galled or non-galled for each ecotype at each adult input. **(F)** Larval phenotyping workflow for quantifying insect performance within galls. Larvae were extracted from individual galled inflorescences, mounted in multi-well plates, imaged and segmented to quantify larval number and body area. **(G)** Gall induction rate in Col-0 and Ler-0 plants following clip-cage no-choice assay at 7 DPO. Bars show pooled gall induction (%) after infestation with 5M+5F adults. Error bars indicate ± binomial SE. Coloured points represent 6 independent paired experiments with Col-0 and Ler-0 assayed side by side; point size indicates total number of plants assessed. *P*-value was obtained by likelihood ratio testing (LRT) of a binomial generalised linear mixed model (GLMM), with experiment as a random effect. **(H)** Larval performance in galled Col-0 and Ler-0 inflorescences at 7 DPO following clip-cage infestation with 5M+5F adults. Larval count, average larval size (area) and summed larval area were quantified for each galled inflorescence. Boxplots show the median and interquartile range, with whiskers extending to 1.5x the interquartile range. Each point represents an individual infested plant; only samples with recovered larvae are shown (Col-0, *n* = 14; Ler-0, n = 18). *P*-values indicate results of Welch’s *t*-tests comparing Col-0 and Ler-0.

To test which host tissues support gall formation, we established a clip-cage infestation assay that confined adults to a defined host region for 24 h (**Fig. 2C**). Neither leaves nor vegetative meristems formed galls after exposure to 5M+5F, whereas actively developing inflorescences at budding-to-early-flowering stages 5.10–6.25 (Boyes *et al*., 2001) did form galls (**Fig. S4C**). We therefore standardised clip-cage infestations to 5M+5F confined to the main inflorescence of *At* plants within this developmental window. Under these conditions, characteristic galls formed in both Col-0 and Ler-0 by 7 DPO (**Fig. 2D**). Using whole-cage and clip-cage assays, we identified actively developing inflorescences as both the main targets of oviposition and reliable tissues for *At* gall induction.

Dissecting the genetic basis of SM-induced gall formation requires reproducible measurements of susceptibility differences between Arabidopsis genotypes. Since galling output reflects the combined effects of insect activity and host response, we used complementary assays to assess: (i) female host choice, (ii) gall induction ability and (iii) larval performance.

Using Col-0 and Ler-0 as representative ecotypes, we first quantified gall induction rate using the clear binary distinction between galled and non-galled inflorescences (**Fig. 2B, D**). Whole-cage choice assays exposed adults to both ecotypes in the same cage, thereby integrating female host choice with gall induction ability. Galls formed exclusively on Ler-0 at lower adult inputs (50–75 adults), while 83% of Ler-0 plants and 33% of Col-0 plants were galled when exposed to a higher adult input (100 adults) (**Fig. 2E**). In clip-cage no-choice assays, Ler-0 again showed a significantly higher gall induction rate than Col-0 (83.1% ± 4.9 versus 62.7% ± 6.3, total percentage ± SE) after accounting for experiment-to-experiment variation (**Fig. 2G**).

To quantify insect performance within individual galls, we established an image-based larval phenotyping workflow. For this, larvae were extracted from individual clip-cage-infested inflorescences, imaged using the natural autofluorescence of the insect cuticle, and segmented to isolate each larva (**Fig. 2F**). At 7 DPO, we used this workflow to quantify larval count, average larval size and total larval load (summed larval area per gall) for each galled inflorescence, with larval load defined as the summed larval area (**Fig. 2H**). Among successfully infested samples from which larvae were recovered, larval number per gall did not differ significantly between Col-0 and Ler-0, whereas average larval size was significantly greater in Ler-0. Total larval load did not differ significantly (**Fig. 2H**).

Together, these assays revealed higher gall induction and larger average larval size in Ler-0 than in Col-0. Our infestation and phenotyping assays provide complementary measures of SM galling output, enabling quantitative comparisons across Arabidopsis genotypes and experimental treatments.

### 3. Tracking gall and larval development reveals the gall-inducing capacity of early larval stages

To define the temporal progression of gall development, we used the clip-cage assay on developmentally matched main inflorescences of early-flowering Arabidopsis. Daily photographs revealed a reproducible progression of symptoms in both Col-0 and Ler-0, with visible galling beginning around 5 DPO and becoming clearly distinguishable by 7 DPO (**Fig. 3A, S5A**). In parallel, we used the larval phenotyping pipeline to resolve daily insect developmental progression following oviposition. Segmented objects were first classified as eggs or larvae based on morphology, and larvae were assigned to L1-L3 using body-area thresholds derived from daily size distributions (**Fig. S5D**). Across 273 Col-0 and Ler-0 infestations sampled from 1–8 DPO, eggs began hatching around 3 DPO, L1 predominated at 3 DPO, L2 at 4 DPO and L3 at 7 DPO, with >95% of larvae classified as L3 by 8 DPO (**Fig. 3D, S5E, F**). The overall developmental sequence was similar between Col-0 and Ler-0 and was consistent with mature L3 larvae beginning to leave Arabidopsis plants from ∼9 DPO (**Fig. 1A**). Thus, visible gall development could be aligned with insect developmental progression: gall symptoms became apparent before L3 predominated and were pronounced by 7 DPO **(Fig. 3A,D)**.

**Fig. 3.**
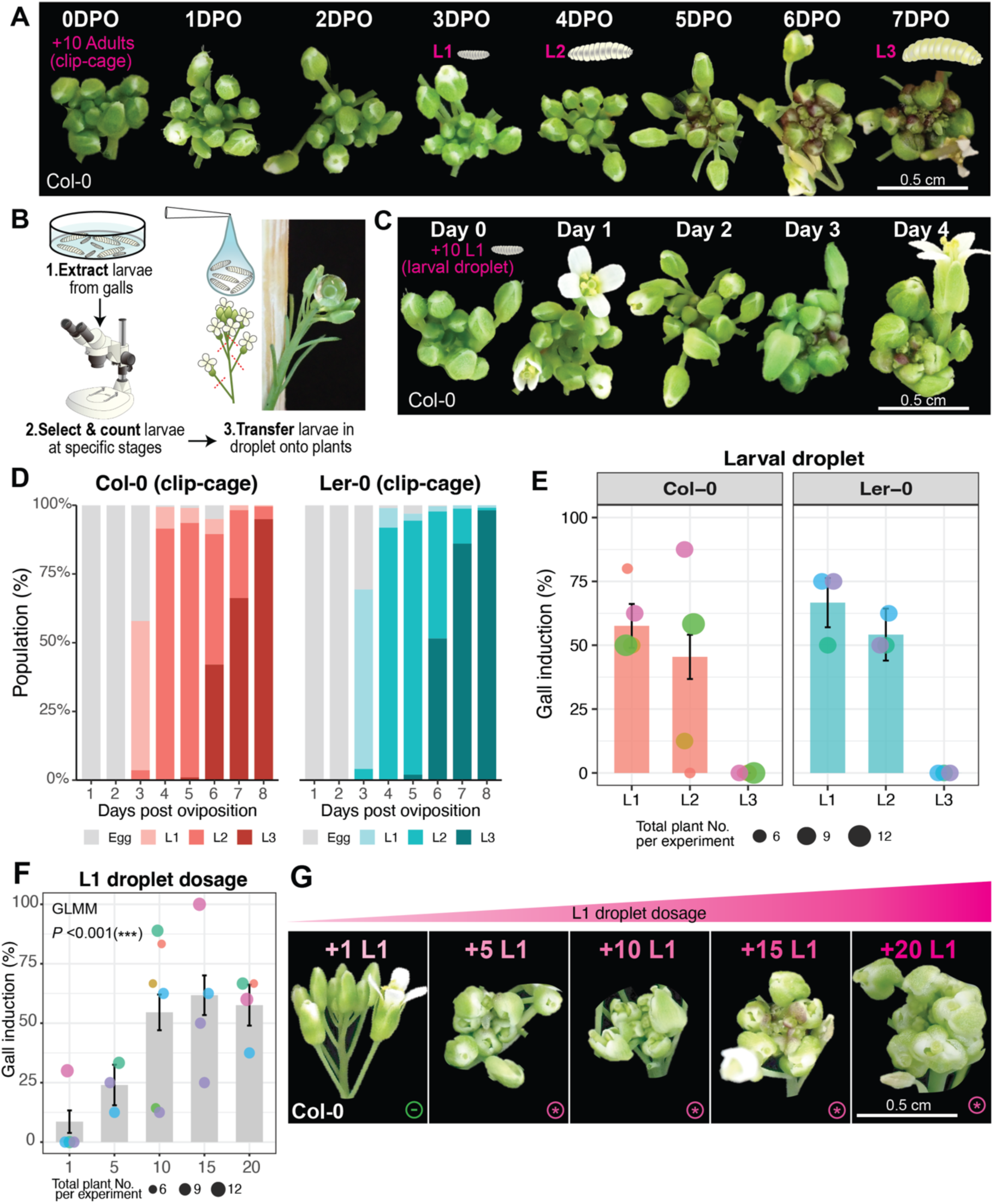
Daily progression of Arabidopsis gall symptoms and Swede Midge larval development following infestation. **(A)** Daily progression of gall phenotypes in Col-0 from 0–7 days post oviposition (DPO) following clip-cage infestation with 5M+5F adults. This experiment was repeated 3 times with similar phenotypes. L1, L2 and L3 indicate the days each larval stage became predominant in the population. Scale bars = 0.5 cm. **(B)** Larval droplet infestation assay workflow. Larvae were extracted from galls, selected at a specific developmental stage (L1, L2 or L3) under a microscope, counted and then transferred onto plant inflorescences in droplets. **(C)** Daily progression of gall phenotypes in Col-0 following application of 10 L1 larvae, shown from day 0–4 after larval droplet application. This experiment was repeated 3 times with similar phenotypes. Scale bars = 0.5 cm. **(D)** Daily progression of egg and larval developmental stages in Col-0 and Ler-0 following clip-cage infestation. Bars show the pooled percentage (%) of the total population assigned to egg, L1, L2 and L3 at each day post oviposition. Data were obtained from 94/170 Col-0 and 97/103 Ler-0 infested plants with positive larval or egg counts, with 10–15 positive plant replicates per day for each genotype (see also Table S1). Eggs are shown in grey, with L1–L3 shown as progressively darker genotype-specific shades. Larval stages were classified by area cutoffs as in **Fig. S5D**. **(E)** Gall induction (%) following application of 5 L1, L2, or L3 larvae to Col-0 or Ler-0 inflorescences, scored 4 days after larval application. Bars show pooled percentage ± binomial SE. Coloured points represent individual experiments; point size indicates the number of plants assessed per experiment. Each plant received all three larval stage treatments on separate inflorescences, enabling matched within-plant comparison (Col-0, *n* = 33; Ler-0, *n* = 24). **(F)** Gall induction (%) following application of 1, 5, 10, 15 or 20 L1 larvae to Col-0 inflorescences, recorded 4 days after larval application. Bars show pooled percentage ± binomial SE. Coloured points represent individual experiments; point size indicates plant number assessed per experiment. Within each experiment, the same plants received the L1 doses tested. The overall dose effect was tested by likelihood ratio testing (LRT) of a binomial GLMM, with experiment as a random effect (*P* = 2.96 × 10⁻⁸). **(G)** Representative gall phenotypes of Arabidopsis (Col-0) inflorescences infested with 1, 5, 10, 15, and 20 L1 larvae. Scale bar = 0.5 cm.

Despite this reproducible population-level progression, individual infestations collected at the same DPO varied in both insect abundance and larval-stage composition, particularly at later time points (**Fig. S5E, F; Table S1**). Thus, DPO provides a useful temporal reference at the population level but does not fully define insect developmental state within individual infestations.

Developmental-stage tracking further clarified the Col-0/Ler-0 susceptibility differences detected following adult infestation. Although total larval number at 7 DPO was similar between ecotypes (**Fig. 2H**), Ler-0 contained a higher proportion of L3 larvae than Col-0 (86.2% versus 66.2%; **Fig. 3D**). This indicates more advanced larval development in Ler-0 at 7 DPO, coinciding with the larger average larval size observed in Ler-0 (**Fig. 2H, 3D**). Differences between ecotypes were also apparent earlier in the interaction. Following adult exposure, Ler-0 plants were more frequently egg-positive at 1–2 DPO than Col-0 plants (89.3% versus 51.2%), although this was not accompanied by higher egg numbers per egg-positive plant (**Table S2**). This was consistent with the whole-cage choice assay, in which galls formed exclusively on Ler-0 at lower adult inputs (**Fig. 2E**). These results indicate that galling outcomes in Col-0 and Ler-0 differ at multiple stages of the interaction, from oviposition to subsequent larval development and gall formation.

To test directly which larval stages were competent to induce galls, we developed a larval droplet assay in which defined numbers of stage-selected larvae were transferred directly onto young inflorescences, bypassing adult oviposition and egg hatching **(Fig. 3B)**. L1 and L2 larvae induced galls in both Col-0 and Ler-0, whereas L3 larvae did not induce galls in either ecotype (**Fig. 3E**). L1 application produced characteristic gall phenotypes, whereas L2 application produced galls accompanied by tissue damage, and L3 produced damage without gall induction (**Fig. S5B**).

To establish a standardised gall-inducing input, we tested how L1 number affected gall induction. Gall induction rate increased significantly with L1 input, rising strongly up to 10 L1 before plateauing at 15–20 L1 (**Fig. 3F**). We observed reduced retention of larger droplets on the inflorescence, which likely contributed to this plateau. We therefore selected 10 L1 as a standardised input, which reproducibly generated the characteristic inflorescence gall phenotype within four days (**Fig. 3C, G, S5C**). Since L1 larvae normally predominated around 3 DPO following adult infestation, the four-day droplet phenotype corresponded closely in timing to the ∼7 DPO phenotype observed after clip-cage infestation (**Fig. 3A, C**). Fixing both larval stage and number with the droplet assay therefore provided a defined insect input and biological time point zero for subsequent analyses of gall development.

These assays link gall progression with insect developmental state and identify L1 and L2 as gall-inducing stages. They resolve gall formation as a coupled host-insect developmental process while defining galling output at several levels: binary gall induction, temporal progression of visible symptoms and quantitative measures of insect abundance and developmental state. This allows galling outcomes to be compared across genotypes and treatments while accounting for insect performance.

### 4. Galling enlarges floral buds and the inflorescence meristem while altering floral development

To define the effects of SM infestation on floral development, we combined a defined larval input with a phyllotactic dissection and positional annotation workflow **(Fig. 4A)**. Floral buds within a single Arabidopsis inflorescence span multiple rapidly changing developmental states, and galling itself also alters floral development **(Fig. 1B, S3A, B)**. To account for this variation, this workflow assigns each bud two positional coordinates, approximating developmental stage and age, to compare galled and non-galled tissues.

**Fig. 4.**
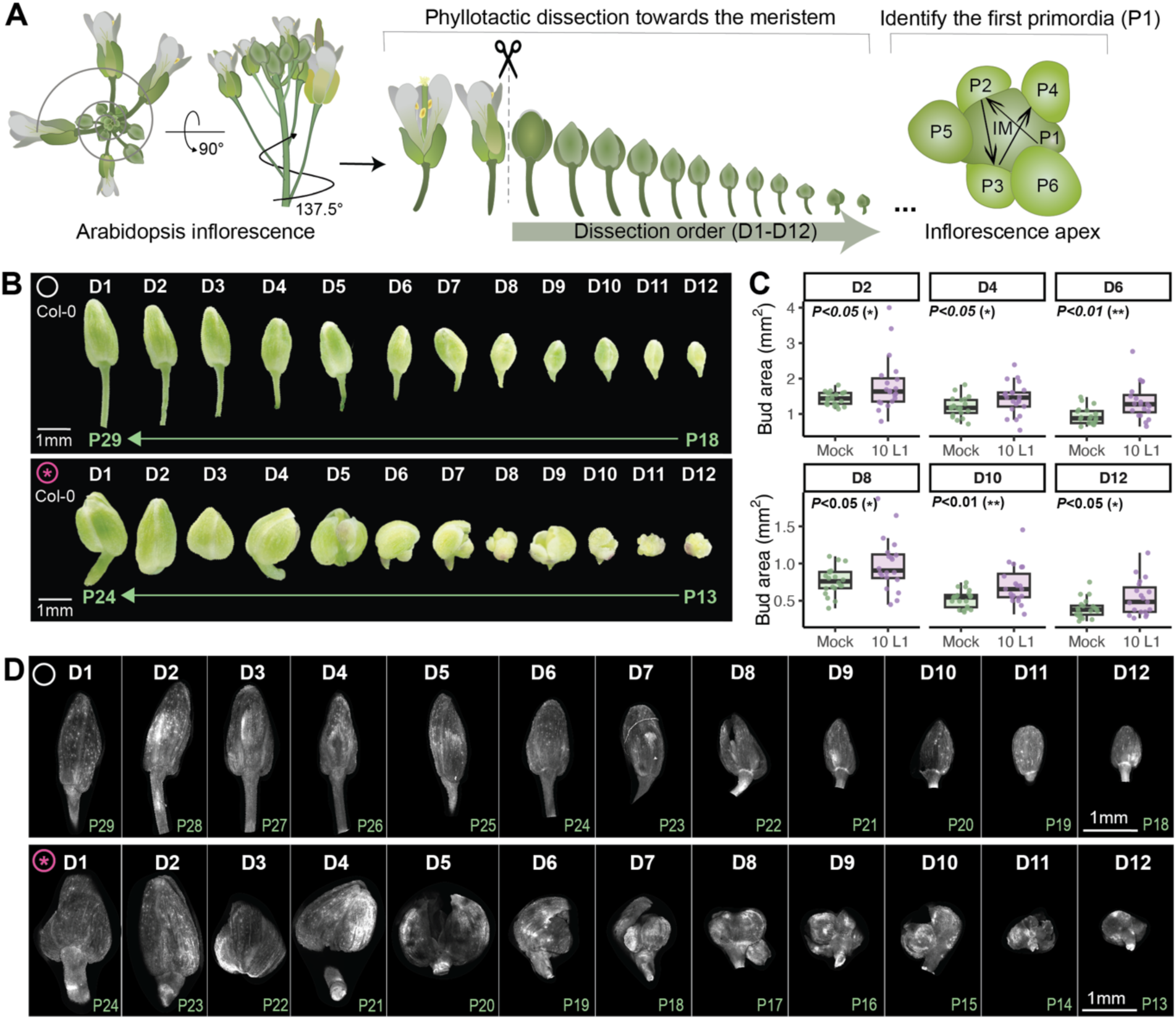
Phyllotactic dissection and quantitative analysis of gall-associated floral bud morphology. Col-0 inflorescences were mock-treated or infested with 10 L1 using larval droplet assays and dissected 4 days after application. White circles indicate mock-treated samples; pink stars indicate infested and galled samples. **(A)** Inflorescence dissection and developmental positional annotation workflow. Floral buds were dissected sequentially along the phyllotactic series towards the inflorescence apex and assigned dissection orders D1–D12. D1 was defined as the first unopened floral bud remaining after removal of all open flowers and late-stage buds from the inflorescence (stage >13). Primordia numbers were assigned by sequential counting from the youngest stage 2 floral primordium (P1) adjacent to the inflorescence meristem (IM) through progressively older primordia and buds (Smyth *et al*., 1990). **(B)** Representative photos of freshly dissected D1–D12 buds from mock-treated and galled inflorescences. Corresponding primordia numbers are shown in green. Scale bars = 1 mm. **(C)** Bud area was quantified from photographs of freshly dissected buds from mock-treated and 10 L1-treated samples at D2, D4, D6, D8, D10 and D12 (see raw images in Fig. S6). Each dissection order included 20–21 mock-treated plants and 19–21 10 L1-treated plants. Points represent individual buds; boxplots show the median and interquartile range. *P*-values are from Welch’s *t*-tests comparing mock and 10 L1 within each dissection order. **(D)** Spinning-disk microscopy images of the same D1–D12 buds shown in (B), displayed as maximum-intensity z-projections. Grey intensity indicates Calcofluor White-stained cell walls. Scale bars = 1 mm.

Buds were dissected sequentially along the phyllotactic series towards the inflorescence apex and labelled D1-D12 according to dissection order, with increasing D number corresponding to progressively younger buds **(Fig. 4A)**. Within the inflorescence apex, P1 was defined as the youngest visible stage-2 floral primordium that had become morphologically distinct from the inflorescence meristem, with successively older primordia numbered P2, P3 and onwards **(Fig. 4A)**. The P series was then traced through the dissected buds, allowing each sampled bud to be assigned both a dissection order and a corresponding P position **(Fig. 4A,B)**.

Four days after application of 10 L1 larvae, freshly dissected buds from galled inflorescences were enlarged and rounder than mock-treated buds at equivalent D numbers **(Fig. 4B, S6)**. Quantification confirmed increased bud area at multiple positions along the series **(Fig. 4C)** and a reduced aspect ratio across the majority of the series **(Fig. S7D)**. Bud area and shape therefore provide quantitative measures of gall morphology in addition to binary gall/non-gall scoring at the inflorescence level.

The dissection workflow also allowed individual buds and the remaining inflorescence apex to be collected while retaining the D and P identities of each sampled bud **(Fig. 4A, 5)**. Dissected material could then be processed for whole-mount imaging and sectioned for histology, revealing tissue organisation and internal floral organs that were not apparent in fresh-tissue photographs **(Fig. 4D, 6, S7A-C)**. This workflow provided position-resolved material for subsequent analysis of gall-associated developmental and cellular changes.

Across 367 buds from 36 infested inflorescences, galling showed a positional bias towards younger buds. Galled buds occurred preferentially at younger positions closer to the inflorescence meristem, as reflected by lower P positions and higher dissection orders **(Fig. S7E, F)**, revealing that younger floral tissues near the meristem were more frequently galled.

To determine whether these changes extended to the inflorescence apex, we examined the meristem and surrounding floral primordia by whole-mount live imaging. The galled apex was notably enlarged, including both the inflorescence meristem and surrounding floral primordia **(Fig. 5A-C)**. Inflorescence meristem area was significantly increased following 10 L1 infestation **(Fig. 5D)**.

**Fig. 5.**
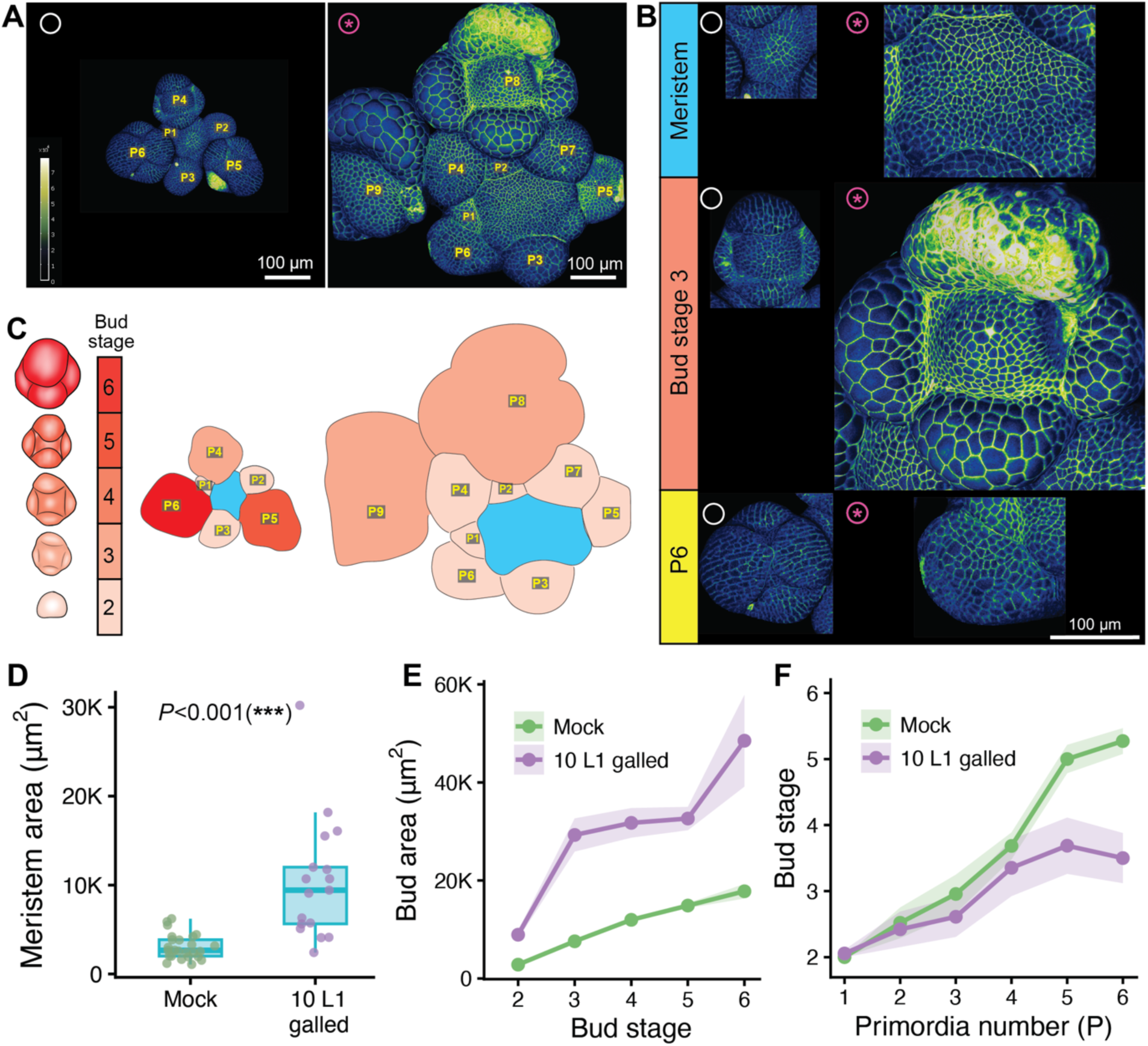
Gall-associated changes in inflorescence meristem and floral primordium development. Col-0 inflorescences were mock-treated or infested with 10 L1 using larval droplet assays and dissected 4 days after application. White circles indicate mock-treated samples; pink stars indicate infested and galled samples. **(A)** Confocal microscopy images of representative mock-treated and galled inflorescence apices, projected onto curved mesh surfaces. Numbers indicate primordia numbers (P); the colour bar indicates propidium iodide staining intensity (arbitrary units). Scale bars = 100 µm. **(B)** Representative meristem and floral primordia from mock-treated and 10 L1-infested apices compared at the same developmental stage (stage 3) or at the same primordia position (P6). Scale bar = 100 µm. **(C)** Schematic demonstration of bud stage assignment across the primordia series shown in (A). Red colour gradient indicates bud stage progression; blue colour indicates meristem. **(D)** Boxplots of inflorescence meristem area from mock-treated and 10 L1 infested apices. **(E)** Mean bud area at each bud developmental stage, shaded area indicates ± SE. Sample sizes: Mock *n* = 121, 10 L1 galled *n* = 109. **(F)** Mean bud stage across primordia series (P1-P6), shaded area indicates ± SE. Sample sizes: Mock *n* = 125, 10 L1 galled *n* = 130.

To distinguish changes in floral primordium size from changes in developmental progression, we compared mock-treated and galled primordia using developmental stage and P position as complementary references. Here, we assigned bud developmental stages based on morphological characteristics of floral organs **(Fig. 5C)** (Smyth *et al*., 1990). When matched by developmental stage, galled primordia or young buds were consistently larger than mock-treated buds across stages 2–6 **(Fig. 5E)**, as illustrated by the stage-3 comparison **(Fig. 5B)**. When matched by P position, buds in galled inflorescences were at earlier developmental stages than mock-treated buds across P1-P6 **(Fig. 5F)**. The representative P6 images show this same difference **(Fig. 5B)**.

This stage difference is unlikely to reflect simply having more primordia, as total primordium counts were comparable between mock-treated and galled apices **(Fig. S7G)**, supporting a delayed bud stage progression in the galled apex. At equivalent P positions, galled buds were also larger **(Fig. S7H)**, showing that they were enlarged despite being developmentally less advanced. Altogether, these results show that 10 L1-induced galling enlarges the inflorescence meristem, floral primordia and young buds, and support delayed progression through normal floral developmental stages.

### 5. Gall formation is associated with altered cell morphology and tissue organisation in floral organs

Having identified pronounced changes in floral-organ growth and development, we next examined the cellular- and tissue-level features associated with gall morphology in Arabidopsis. Whole-mount spinning-disk and confocal imaging, scanning electron microscopy and histological sections revealed cellular changes in galled sepals and pedicels **(Fig. 6, S8)**.

**Fig. 6.**
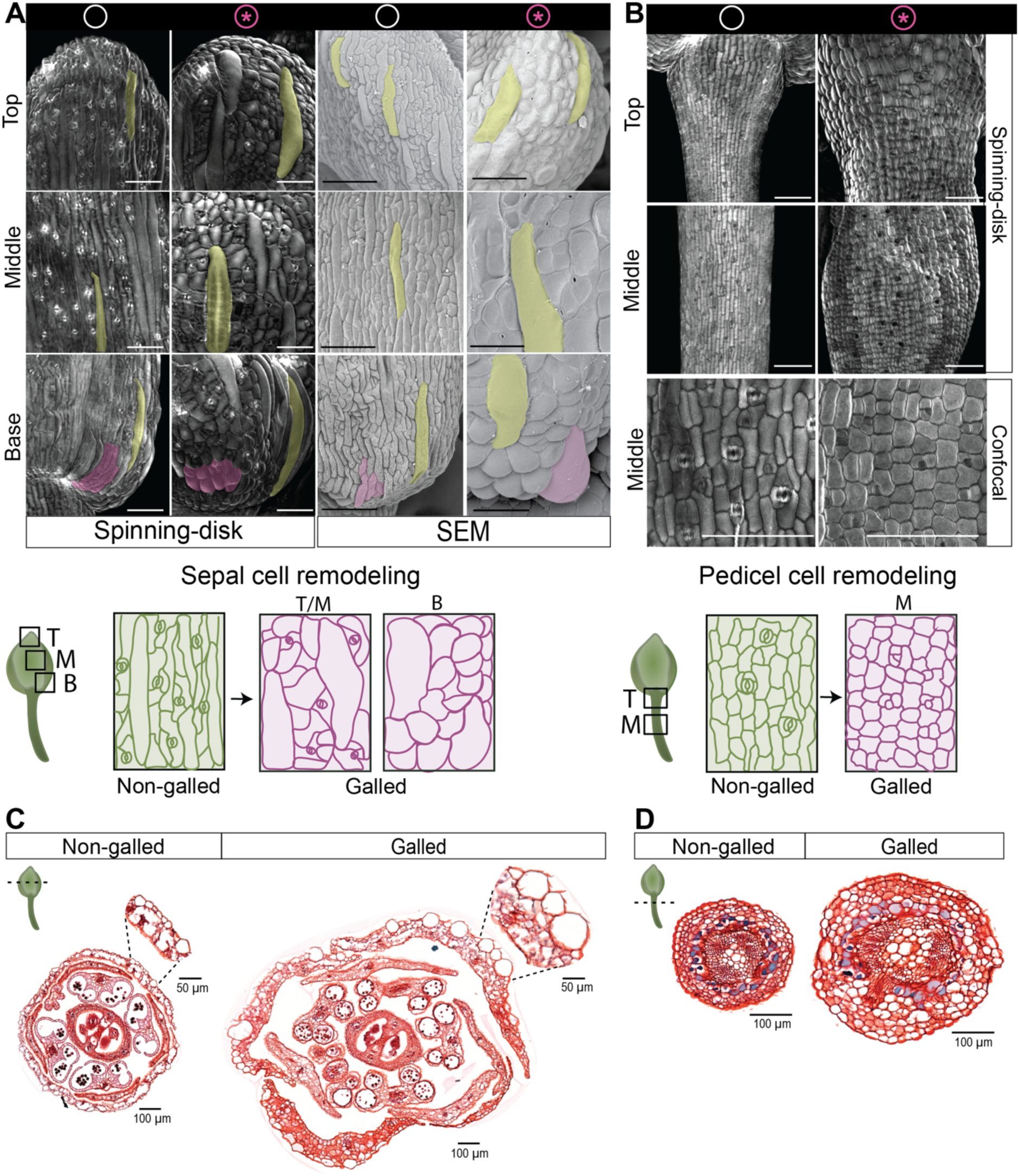
Gall-associated cellular changes in floral buds and pedicels. Col-0 inflorescences were mock-treated or infested with 10 L1 larvae using larval droplet assays and analysed 4 days after application. White circles indicate mock-treated samples; pink stars indicate infested and galled samples. **(A)** Representative spinning-disk and SEM images of cellular changes in non-galled and galled sepals. Giant cells (Roeder *et al*., 2010) in the galled and non-galled sepal epidermis are pseudocoloured green. Gall-associated “bubbly” cells at the sepal base and comparable small pavement cells in non-galled sepals are pseudocoloured purple. Spinning-disk images are displayed as maximum-intensity z-projections; grey signal indicates Calcofluor White-stained cell walls. Scale bars = 100 µm. The schematic directly below illustrates cellular patterns in non-galled and galled sepals. Top (T), Middle (M), and Base (B) indicate the corresponding sepal regions in the microscopy images above. **(B)** Representative spinning-disk and confocal microscopy images showing cellular changes in non-galled and galled pedicels. Images are displayed as maximum-intensity z-projections; grey signal indicates Calcofluor White-stained cell walls. Scale bars = 100 µm. The schematic directly below illustrates cellular patterns in non-galled and galled pedicels. Top (T) and Middle (M) indicate the sepal–pedicel junction and middle pedicel region, respectively, corresponding to the microscopy images above. **(C)** Transverse histological sections of non-galled and galled floral buds stained with Safranin O and Fast Green. Magnified views show sepal cellular morphology, including enlarged cells in the galled bud. Scale bars = 100 µm for whole-bud sections and 50 µm for magnified views. **(D)** Transverse histological sections of non-galled and galled pedicels stained with Safranin O and Fast Green. Scale bars = 100 µm.

In non-galled sepals, epidermal cells showed the characteristic pattern of elongated pavement cells interspersed with giant cells (Roeder *et al*., 2010). In galled sepals, epidermal cells were enlarged across the top, middle and base of the sepal; giant cells were further enlarged and less elongated **(Fig. 6A)**. Towards the sepal base, surrounding pavement cells also became enlarged and rounded, often forming clusters that gave the surface a characteristic “bubbly” appearance **(Fig. 6A)**. These bubbly cells were visible in freshly dissected galled buds, where they appeared transparent and inflated **(Fig. S8C)**. Transverse histological sections further showed that galled sepals were thicker, with cell enlargement also evident in internal tissues beyond the epidermis **(Fig. 6C, S8A)**.

At the whole-bud level, histological sections revealed extensive changes in internal tissue organisation. Galled buds were enlarged, with thicker sepals and petals as well as expanded spaces between floral organs evident in sections from the top, middle and base of the bud **(Fig. 6C, S8A)**. Swede Midge larvae were observed within the expanded spaces between floral organs (**Fig. S8A**).

Galled pedicels also showed changes at both the epidermal and internal tissue levels. Compared with the elongated epidermal cells of non-galled pedicels, cells in galled pedicels were shorter and broader, both near the sepal-pedicel junction and further along the pedicel (**Fig. 6B**). Histological sections showed that galled pedicels were thicker than non-galled controls, with expanded internal tissues and altered tissue organisation (**Fig. 6D, S8B**). Thus, pedicel thickening was associated with changes in both epidermal cell morphology and internal tissue organisation.

Gall-associated changes in cell morphology and tissue organisation across sepals, petals and pedicels provide reproducible cellular and histological readouts of SM-induced gall formation.

## Discussion

### A tractable model for insect gall research

Mechanistic understanding of insect gall formation remains limited by the scarcity of experimental systems that combine genetic accessibility with precise control of insect and host development (Giron *et al*., 2016). Gall formation reflects the interaction of two dynamic inputs, insect-derived signals and host responses, and resolving their contributions requires control over developmental variation in both organisms. Existing systems, including Hessian Fly–wheat (Stuart *et al*., 2012), gall weevil–dodder (Ushima *et al*., 2024) and *Hormaphis cornu*–Witch Hazel (Korgaonkar *et al*., 2021), solve parts of these problems, but none combines controlled gall induction, developmental tracking of both partners and the molecular resources of a model host. Here, we establish the Arabidopsis-SM model as a resource that brings these capabilities together, establishing an experimentally tractable system for insect gall research.

We established that SM completes its life cycle on Arabidopsis and induces characteristic inflorescence galls resembling those on Brassicaceae crops. Using direct larval application, we identified L1 and L2, but not L3, as gall-inducing stages. In the host, we found that galling enlarged floral buds, floral primordia and the inflorescence meristem and delayed progression through floral stages. We also identified characteristic changes in cell morphology and tissue organisation that provide additional readouts of gall formation. These findings establish a framework for defining galling outcomes while controlling and accounting for developmental variation in both partners.

### Controlling and accounting for insect developmental variation

Given that gall induction and progression depend on the insect activity, monitoring insect abundance and developmental state is essential. Our analysis showed that individual infestations collected at the same DPO differed in both larval abundance and developmental-stage composition, even when adult input and host developmental stage were standardised **(Fig. S5E, F, Table S1)**. Our semi-automated larval phenotyping pipeline enables quantitative monitoring of these parameters across individual infestations and time points, providing a scalable approach to account for insect developmental variation. The Col-0/Ler-0 comparison illustrates its value: although larval numbers did not differ significantly between ecotypes among samples with recovered larvae, the higher proportion of L3 larvae and larger average larval size in Ler-0 at 7 DPO indicated more advanced insect development **(Fig. 2H, 3D)**.

While monitoring allows us to account for developmental variation, controlling insect input can reduce this variation experimentally. Adult infestation introduces variation in egg number and the timing of oviposition and hatching. Our larval droplet assay bypasses these processes by applying defined numbers of stage-selected larvae directly to host inflorescences, establishing a controlled time zero for plant-insect interactions.

Standardised application of 10 L1 larvae produced characteristic galls within four days, providing a short and defined timeline for molecular and phenotypic analyses **(Fig. 3C)**. Nevertheless, fixing the initial larval stage and number does not eliminate subsequent variation in insect development or galling success. Importantly, the ability to isolate living larvae and reapply them to host tissues also provides opportunities for direct insect manipulation. Larvae could be treated with dsRNA or other reagents before transfer to Arabidopsis, potentially enabling causal tests of candidate insect gene function through effects on larval development and gall formation.

### Controlling and resolving host developmental variation

Our data demonstrate that galling outcomes vary with host developmental stage and tissue type. Maintaining consistent environmental conditions, including humidity, is also important for reproducible infestation. To reduce host developmental variation, we synchronised flowering through a short-day to long-day transition, selected plants at comparable developmental stages (Boyes *et al*., 2001) and standardised infestation to actively developing inflorescences. Progressing from whole-cage to clip-cage and larval droplet assays further improved control over the target tissue and initial infestation conditions.

Even when plants are developmentally matched at infestation, individual floral buds within a single inflorescence vary substantially in developmental stage. Galling itself alters normal floral development, making it difficult to identify developmentally equivalent controls based on bud position or morphology alone. Our phyllotactic dissection and positional annotation workflow provides complementary coordinates for comparing galled and non-galled tissues **(Fig. 4A)**. Primordium position (P) locates individual primordia relative to the inflorescence meristem, while dissection order (D) provides a reproducible sampling series from older to younger buds. However, equivalent positional coordinates do not necessarily indicate equivalent developmental stages. Combining positional information with direct floral staging therefore provides an informative basis for comparison. Using this approach, we observed enlarged primordia and young buds in galled inflorescence apices despite less advanced floral developmental stages (**Fig. 5, S7H**).

### Defining and interpreting galling output

Galling is a complex trait that cannot be represented by a single measure. Thus, it requires complementary measures to assess gall induction, morphology and developmental progression. Whole-cage choice assays integrate adult host preference and subsequent infestation success, while clip-cage assays standardise adult exposure to defined host tissues. Binary gall scoring at 7–8 DPO quantifies gall induction frequency across genotypes and treatments. Our developmental and cellular analyses provide additional measures of gall formation across multiple scales, from the inflorescence apex to individual cells in differentiated floral organs. Meristem and primordium enlargement altered floral stage progression, and changes in cell size, shape and tissue organisation provide distinct readouts of the host response **(Fig. 4–6)**. Gall formation was concentrated in younger floral tissues near the meristem, where developmental changes were also apparent (**Fig. 5, S7E, F**). These observations suggest greater gall competence in younger, less differentiated floral tissues and show that SM-associated changes extend to early floral development. However, the precise tissue or cell type in which gall induction begins remains unresolved. Greater spatial and temporal resolution is therefore needed to determine when and where host development first diverges during gall formation and to identify insect-plant contact sites.

Insect performance provides a complementary functional outcome of the host–insect interaction rather than a direct measure of gall severity. Larval number, average size, developmental-stage composition and total larval load capture different aspects of insect establishment and development, providing quantitative measures of insect performance before visible gall symptoms emerge. Galling notably disrupts normal floral development, making host developmental stage alone insufficient to describe the progression of the interaction. Tracking insect development alongside host morphological and cellular changes provides an additional reference for the progression of the interaction. Relating these measurements will allow us to investigate how specific host developmental changes are associated with insect performance and whether genetic or molecular perturbations affect gall formation, insect development, or both.

### Towards mechanistic analysis of insect gall formation

Our finding that L1 and L2, but not L3, larvae induce galls identifies the early larval stages as the gall-inducing period and demonstrates that oviposition or ovipositor-mediated interaction with host tissue is not required for gall induction (**Fig. 3E, S5B**). This provides a biological basis for prioritising candidate insect effectors using available SM salivary-gland transcriptomic resources, particularly comparisons between L1 and L3 larvae (Mori *et al*., 2023, 2024). Arabidopsis genetics offer an established route for testing the contributions of host genes to gall formation, while developing genetic manipulation, gene-silencing or other genetic approaches in SM would be an important next step towards testing the causal roles of candidate insect effectors.

The Arabidopsis–SM model now provides a framework for investigating how insect-derived inputs and host developmental state contribute to defined galling outcomes, laying the foundation for future mechanistic studies of insect-induced gall formation.

## Materials and Methods

### 1. Plant materials and growth conditions

*Brassica napus* cultivars Westar (IPK accession number 1052), Drakkar (IPK accession number CR 3514) and AC Excel were used to rear Swede Midge under controlled laboratory conditions. Seeds were germinated on vermiculite for one week, and two seedlings per pot were transplanted into 16 cm pots containing commercial substrate (Gramaflor® Topf-TopXL, Germany). Plants were grown under long-day (LD) conditions (16 h light / 8 h dark) at 21 °C and a photosynthetic photon flux density (PPFD) of 166 µmol m⁻² s⁻¹. After approximately 6-7 weeks, plants were transferred to a climatic chamber under the same photoperiod and light conditions, and at ∼75% relative humidity for infestation experiments.

Rapid Cycling *Brassica rapa* plants (Wisconsin Fast Plants®, Standard Variety seeds) were grown and infested at 22 °C during the day and 20 °C at night under LD conditions and at ∼70% relative humidity.

Arabidopsis ecotypes Col-0 and Ler-0 were grown in a 4:1 mixture of commercial soil (Substrat 2 + GreenFibre®, Klasmann-dellmann) and perlite in pots measuring 6.5 x 6.5 x 5 cm or 4.1 x 4.5 x 5 cm. Before sowing, seeds were stratified for 3-4 days in MilliQ water at 4 °C in the dark to synchronise germination. Plants were grown for 4 weeks under short-day (SD) conditions (8 h light / 16 h dark) then transferred to LD conditions to induce flowering. Both SD and LD chambers were maintained at 21 °C and a PPFD of 166 µmol m⁻² s⁻¹. *At* plants at early flowering stages (Boyes *et al*., 2001) were transferred to a climatic chamber under the same LD condition and light settings, and at 75% relative humidity for infestation experiments.

### 2. Swede Midge colony rearing and maintenance

The SM colony was established from natural populations infesting *B. napus* and *B. oleracea* collected from agricultural fields in Switzerland and Germany. The colony was maintained on *B. napus* cultivars Drakkar, Westar and AC Excel at the Gregor Mendel Institute of Molecular Plant Biology (GMI). Two pots of *B. napus* at BBCH stages 57-61(LANCASHIRE *et al*., 1991) were placed in a rearing cage (BugDorm-BDE4590, 47 x 47 x 93 cm) with 800-2000 newly emerged adult SM collected from the emergence cages. To set up the emergence cages, infested *B. napus* plants with galls were trimmed 7 days after oviposition (DPO), and the trimmed pieces were placed on fresh, moistened soil in cages (BugDorm-4F3030, 25 x 25 x 25 cm). The next generation of adults emerged ∼10-15 days later. Details of rearing at the University of Alberta were previously described (Lopez *et al*., 2026).

### 3. Life cycle and adult emergence tracking

To characterise SM life cycle timing on different hosts, whole-cage no-choice experiments were performed using two plants of the same host per oviposition cage (BugDorm-4F4545, 47.5 x 47.5 x 47.5 cm; or BugDorm-1, 30 x 30 x 30 cm MegaView Science, Taiwan). Hosts were Arabidopsis Col-0 at stages 6.10-6.20, 10-20% flowers open (Boyes *et al*., 2001), *B. napus* cultivar Westar and Drakkar at BBCH stages 57-59 (LANCASHIRE *et al*., 1991)(stage 57: individual floral buds visible but still closed), and Rapid Cycling *B. rapa* with individual floral buds visible but still closed. Each cage received 100 adult SM (50 males and 50 females, 50M+50F), with no restriction on oviposition duration. At 7 DPO, galled material was trimmed from the plants and placed on moist soil. Pots were sprayed daily with water. Adult emergence was monitored daily until no adults emerged for three consecutive days **(Fig. S2)**.

Photoperiod and temperature influence diapause entry in SM, with warm, LD conditions associated with reduced diapause frequency (Marteaux *et al*., 2012). Diapause frequency was not quantified in the present study.

### 4. Pupal collection and imaging

Commercial substrate was sieved (0.5 mm mesh size), placed in a 4.1 x 4.5 x 5 cm container and moistened with 10 mL of water. Galled *B. napus* inflorescences were trimmed at 7 DPO and placed on the moist soil surface. The container was watered daily with 5 mL of water. After 9 days, the soil was washed with deionised water through a 0.5 mm sieve. The retained material was transferred gradually into 50 mL falcon tubes, which were then filled to 50 mL with deionised water. Floating material was collected in 6 cm petri dishes. Cocoons were picked out with forceps, kept on damp filter paper, and opened with dissecting pins under a stereomicroscope to extract the pupae.

For imaging, pupae were fixed in 70% ethanol for 2 h and imaged using a Zeiss AxioZoom.V16 (PlanNeoFluar Z 1.0x objective, NA 0.25, 5x zoom). Images were acquired and exported using Zeiss ZEN software (version 3.7.97).

### 5. Swede Midge infestation protocols

#### Whole-cage choice and no-choice assays

For whole-cage no-choice assays, two plants of the same Arabidopsis ecotype were placed in an infestation cage (BugDorm-4F3030). For whole-cage choice assays, one Col-0 and one Ler-0 plant were placed in each cage. Adult SM were collected from several emergence cages, mixed and transferred to the cages for infestation. For no-choice assays, 100 adults (50M + 50F) were used per cage; for choice assays, increasing numbers of adults were used, with no control of female to male ratio.

#### Clip-cage no-choice assays

The main inflorescence was enclosed in a clip-cage (25 mm in diameter; Entomopraxis, G4150), assembled around the stem and secured with clips. The cage was supported with bamboo sticks to keep the inflorescence upright (**Fig. 2C)**. Newly emerged adults (<24 h old) were collected from emergence cages using a mouth aspirator (S-type, 6 mm; Entomopraxis, D603). 5M + 5F were transferred to each clip-cage, which was then closed with a stopper. Adults and clip-cages were removed after 24 h to end the oviposition period. Control inflorescences were caged without adults.

#### Larval droplet assay

Inflorescences of infested *B. napus* were collected at 3 DPO for L1 larvae, 4 DPO for L2 larvae, and 7 DPO for L3 larvae. Larvae were extracted by submerging inflorescences in 1xPBS supplemented with 0.001 % (v/v) Silwet and incubated in 5 mL Eppendorf tubes for 2 minutes with gentle agitation. The suspension was transferred to a small petri dish and inspected under a stereomicroscope (Zeiss Stemi 2000). Defined number of larvae were pipetted in 10-15 μl droplets directly onto Arabidopsis inflorescences after removal of all open flowers **(Fig. 3B)**. Plants were transferred to rearing cages 30 min after application to allow the droplets to settle. Mock-treated inflorescences received the same volume of extraction solution without larvae.

### 6. Gall assessment and larval phenotyping

To monitor gall development, main inflorescences were photographed before infestation (0 DPO or Day 0) and daily for 4 days after larval droplet application, or for 7 days after the clip-infestation. Unless otherwise specified, galling phenotypes were assessed at 8 DPO for whole-cage no choice assay, 7 DPO for whole-cage choice assay and clip-cage no choice assay, and 4 days after larval application.

#### Larval phenotyping workffow

Egg and larval populations were monitored after clip-cage infestations of Col-0 and Ler-0 plants. Inflorescences were collected in 25% (v/v) ethanol at 1-10 DPO and incubated at 4 °C for 24 h. The suspension containing eggs and larvae was transferred to a 48-well plate and imaged using a CellDiscoverer 7 widefield imaging system (Zeiss) **(Fig. 2F)**. Segmentation used the natural autofluorescence of the insect cuticle, acquired in the DAPI channel (emission 450–470 nm). Images were processed in Fiji (ImageJ 1.54) (Schindelin *et al*., 2012) using maximum-intensity projections, followed by a gamma correction to reduce uneven background and plate-associated artefacts. The images were then processed in QuPath (Bankhead *et al*., 2017)(v0.7.0) for segmentation, manual correction and quantification. Cellpose-SAM (Pachitariu *et al*., 2025)was trained to generate two specialised models: one for eggs and early-stage larvae and another for later-stage larvae. Initial training annotations and post-segmentation corrections were performed using SAM in QuPath (Kirillov *et al*., 2023; Sugawara, 2023; Burri *et al*., 2026).

#### Larval performance quantification

Larval performance metrics were calculated for each infested inflorescence level at 7 DPO. Total larval count was defined as the number of larvae detected, average larval size as the mean larval body area measured after extraction and imaging, and larval load as the summed larval area per gall. Only plants with positive larval counts were included.

#### Egg and larval developmental-stage tracking

Infested plants were sampled daily and processed using the larval phenotyping workflow. Eggs were classified using visible morphological differences from larvae. Larvae were assigned putative instar stages using body-area thresholds. Morphological features (e.g. spatula presence) corroborated the classification of a subset of L3 larvae; L1 and L2 assignments were based on body area alone. Larval stage thresholds were defined using the combined Col-0 and Ler-0 datasets derived from 4,290 objects sampled at 1-8 and 10 DPO. Egg and larval area distributions were first examined in pooled density plots, and candidate boundaries were identified from valleys between size distributions. These boundaries were then evaluated against the day-by-day area distributions to check whether stage assignments followed the expected temporal progression (**Fig. S5D**). Final cutoffs were defined as L1 <32,000 µm², L2 = 32,000–110,000 µm², and L3 >110,000 µm².

### 7. Inflorescence dissection, imaging and analysis

#### Assignment of dissection order and primordia number to fforal buds

To compare floral buds at equivalent developmental positions between mock-treated and infested inflorescences, each bud was assigned a dissection order and a primordium number **(Fig. 4A)**. Open flowers and late-stage floral buds (stage > 13) (Smyth *et al*., 1990) were removed and the first unopened floral bud remaining was designated D1 (approximately stage 11-12). Subsequent buds were dissected sequentially along the phyllotactic series towards the inflorescence meristem and designated D2–D12. Increasing dissection order therefore corresponded to progressively younger buds.

After D1-D12 buds were collected, additional younger buds were removed until the remaining floral buds, floral primordia and inflorescence meristem could be resolved by microscopy (the imaged inflorescence apex) (**Fig. 5A**). The youngest visible stage-2 (Smyth *et al*., 1990) floral primordium was designated P1, and progressively older primordia and buds were assigned increasing primordia numbers according to their position along the phyllotactic series. The angular position and size of galled buds were used to guide P number assignment. Buds removed between the imaged inflorescence apex and D12 were included in this count, allowing the primordium number of D12 to be determined and the series extended sequentially from D12 to D1.

#### Bud area and aspect ratio quantification

Photographs of freshly dissected floral buds with assigned dissection orders were analysed (**Fig. S6**). Image processing and morphological quantification were performed using Fiji. Each photograph was spatially calibrated using its scale bar. Images were first converted to 8 bit and binarized using Huang’s automated thresholding with manual adjustments to segment the buds from the background. Pedicels were manually excluded from bud outlines, and individual buds were detected using the Analyze Particles function. The projected area of each bud was measured from its segmented outline. In total, 122 buds from mock-treated plants and 120 buds from 10 L1-treated plants were quantified, with 20-21 mock-treated and 19-21 10 L1-treated plants at each dissection order.

Bud shape was quantified using aspect ratio (AR) calculated in Fiji as the major-axis length divided by the minor-axis length of the best-fit ellipse. The major axis represents the longest axis of the fitted ellipse, and the minor axis represents the shortest axis, perpendicular to the major axis. An AR of 1 indicates a perfect circular fitted ellipse, whereas higher values indicate greater elongation. AR is independent of rotation within the image plane but can be affected by out-of-plane tilt.

#### Whole-mount live imaging and meristem and bud area quantification

To image the inflorescence meristem, inflorescence apices remaining after D1–D12 sampling were mounted upright in petri dishes (60 mm x 15 mm; VWR 25384-092) containing 1% agarose in water and further dissected under a Zeiss Stemi 2000-C stereomicroscope to expose floral buds at stage 6 or earlier (Smyth *et al*., 1990). Samples were stained with 0.1 mg/mL propidium iodide (PI) and 0.1% (v/v) Silwet in water for 5 min, rinsed 3–5 times in deionised water and imaged immediately using a Zeiss LSM 800 Upright Examiner Z1 confocal microscope with a 20x Plan-Apochromat water-dipping lens (NA 1.0). Excitation was at 488 nm, with emission collected at 488–650 nm for PI.

To visualise the inflorescence meristem surface, raw confocal CZI files were converted to TIFF stacks in Fiji, and the PI channel was imported into MorphoGraphX (Reuille *et al*., 2015; Strauss *et al*., 2022). Surface meshes were generated using Gaussian Blur with default settings, Edge Detect, and Marching Cubes (5 μm cubes). The mesh surface was refined using Subdivide Mesh and Smooth Mesh functions, repeated 2–3 times. The PI signal was projected onto the surface from depths of 3–6 μm for galled samples and 2–4 μm for mock-treated samples, with the mesh surface defined as 0 μm.

Projected meristem and bud area were measured from maximum-intensity projections of inflorescence apices; each outline was traced with the Freehand Selection tool and the area calculated using Fiji.

#### Whole-mount imaging of fixed and stained fforal buds

D1-D12 buds were fixed individually in a 96-well plate containing 4% paraformaldehyde (PFA; aqueous solution, EM grade, EMS-15714) in 1x PBS. The plates were subjected to 2–3 vacuum cycles of 10-15 min per cycle, then incubated overnight at 4°C. The PFA solution was then removed the following day, and the buds were washed twice with 1x PBS. Samples were cleared in ClearSee solution for 4–5 days until sufficiently transparent for whole-mount imaging (Ursache *et al*., 2018). Cleared samples were stained in ClearSee solution containing 0.1% Calcofluor White (Santa Cruz, 4193-55-9) and 0.2% Basic Fuchsin (Abcam, ab146335) for 2–3 days at room temperature (RT) with gentle agitation on a bench rocker (Benchmark, Benchwaver). Samples were then washed with ClearSee to remove excess stain.

Images were acquired using an inverted Olympus spinning-disk confocal microscope with SoRa super-resolution, equipped with a 10x air objective (NA 0.4). Excitation wavelengths were 405 nm for Calcofluor White and 561 nm for Basic Fuchsin, with emission collected at 410–435 nm and 600–650 nm, respectively. Maximum-intensity projections were generated in Fiji for each channel and displayed separately or merged.

#### Scanning electron microscopy

Scanning electron microscopy (SEM) of plant and insect samples was performed at the Electron Microscopy Service of the Vienna BioCenter. Samples were fixed in 4% PFA in 1× PBS, subjected to three 5-min vacuum pulses, and incubated overnight in fresh fixative at room temperature. Samples were then dehydrated through an ethanol series and critical-point dried using liquid CO₂ (Polaron E300 apparatus). Dried samples were mounted on activated carbon strips and sputter-coated with gold–palladium (4:1) using a Baltec SCD005 sputter coater. Images were acquired on a JEOL JSM-5410 scanning electron microscope using Autobeam software on the ISIS platform (Oxford Instruments), with a scan speed of 200 s per image, an accelerating voltage of 10–15 kV and a working distance of 25 mm.

#### Sample preparation for histology and sectioning

Mock-treated and 10 L1-treated Arabidopsis inflorescences were dissected as described above. Following dissection, D1-D12 buds were kept individually in a 96-well plate in 1x PBS at 4 °C. The P23 bud from each sample was selected and embedded in 2% (w/v) low-melting-point agarose (Lactan, 63515). Agarose blocks were left to solidify at RT, submerged in 4% PFA under vacuum for 30 min, then incubated at 4°C for 2-3 days. Following fixation, samples were washed 2–3 times with 1× PBS prior to paraffin embedding. Dehydration, embedding, sectioning and staining were performed at the Histology Facility of the Vienna BioCenter. Samples were dehydrated overnight using an automated tissue processor (DiaPath Donatello Series 2) then embedded in paraffin at 56 °C using a tissue embedding station (Tissue-Tek TEC, Sakura). Paraffin blocks were sectioned transversely at 10 µm using a microtome (Microm HM 355 S Leica), and sections were collected on glass slides, and dried overnight at 50 °C. Prior to staining, sections were dewaxed using an automated stainer (Gemini AS). Sections were stained with 0.05% Fast Green for 5 min, rinsed in 1% acetic acid for 10 s, and stained with 0.1% Safranin O for 5 min. They were then dehydrated sequentially in 95% and 100% ethanol for 1 min each and air-dried. Coverslips were mounted using a Tissue-Tek GLC coverslipping machine. Sections were scanned, images were examined using QuPath (v0.7.0), and selected views of the sections were exported for visualisation.

### 8. Graphs and Statistics

Graphs and statistical analyses were performed in R (version 4.4.1, R Core Team, 2024).

**Box plots** show the median and interquartile range, with whiskers extending to 1.5x the interquartile range, and individual points represent plants or cages as specified in figure legends. Differences between Col-0 and Ler-0 or between mock-treated and infested samples were assessed using two-sided Welch two-sample *t*-tests, which accommodate unequal variances and unequal sample sizes. Total adult emergence per cage was compared among host species using one-way ANOVA followed by Tukey’s HSD test. Different letters indicate significant pairwise differences at Tukey-adjusted *P* < 0.05.

**Violin plots** show the distribution of observed values, with violin width indicating data density. Individual points represent floral buds or inflorescences, as specified in the figure legends, and black horizontal bars indicate medians. Bud positions were compared using two-sided paired Wilcoxon signed-rank tests on plant-level medians, restricted to plants containing both galled and non-galled buds. Total primordia counts were compared between mock-treated and galled inflorescences using a two-sided Wilcoxon rank-sum test.

**Gall induction (%)** was calculated as 100x the number of galled plants or treated inflorescences divided by the total number assessed, according to the scoring unit used for each assay. Samples were classified with a binary galled/non-galled outcome based on the characteristic galling phenotype at the assessment time specified in section 6. Bars show pooled percentages ± binomial SE. Differences in gall induction between genotypes or treatments were tested using a binomial generalised linear mixed model (GLMM), with independent experiment as a random effect. Significance was assessed using likelihood ratio tests (LRTs), with Tukey-adjusted pairwise comparisons where indicated (*P* < 0.05). Statistical significance was defined as *P* < 0.05 (*), *P*<0.01 (**) and *P*<0.001(***).

## Acknowledgment

We are thankful for the Vienna Biocenter Core Facilities (VBCF): BioOptics Facility for advanced imaging and microscopy support, the Electron Microscopy Facility for ultrastructural analysis and imaging, the Histology Lab, the Plant Sciences Facility and the Child Care Center. We thank Julia Holakovsky, Devika Talwar, Miguel Rodrigez, Kelly Dunfield, Kristen Guelly, Sadey Hamilton, Jaymie Martin and Alexsandra Beatriz Pinheiro for the technical support. Byron Rusnak was funded by the Vienna BioCenter Interdisciplinary Postdoctoral Program 3 (VIP3) that receives funding from the European Union’s Horizon Europe Research and Innovation Programme under the Marie Skłodowska-Curie grant agreement No. 101217025. Boyd Mori was funded by the NSERC Discovery Grant (RGPIN-2021-02479) and by the NSERC Industrial Research Chair (545088) and partner organizations (Alberta Grains, Alberta Canola Producers Commission, Alberta Pulse Growers Commission). We thank Silvia Ramundo, Ortrun Mittelstein Scheid and Liam Dolan for the critical reading of the manuscript. For open access, the author has applied a CC BY public copyright license to any Author Accepted Manuscript version arising from this submission. Open access funding provided by Gregor Mendel Institut fur Molekulare Pflanzenbiologie GmbH.

