## Supplemental Figures and Legends for "Arabidopsis–Swede Midge as an experimentally tractable insect gall model to dissect plant reprogramming"

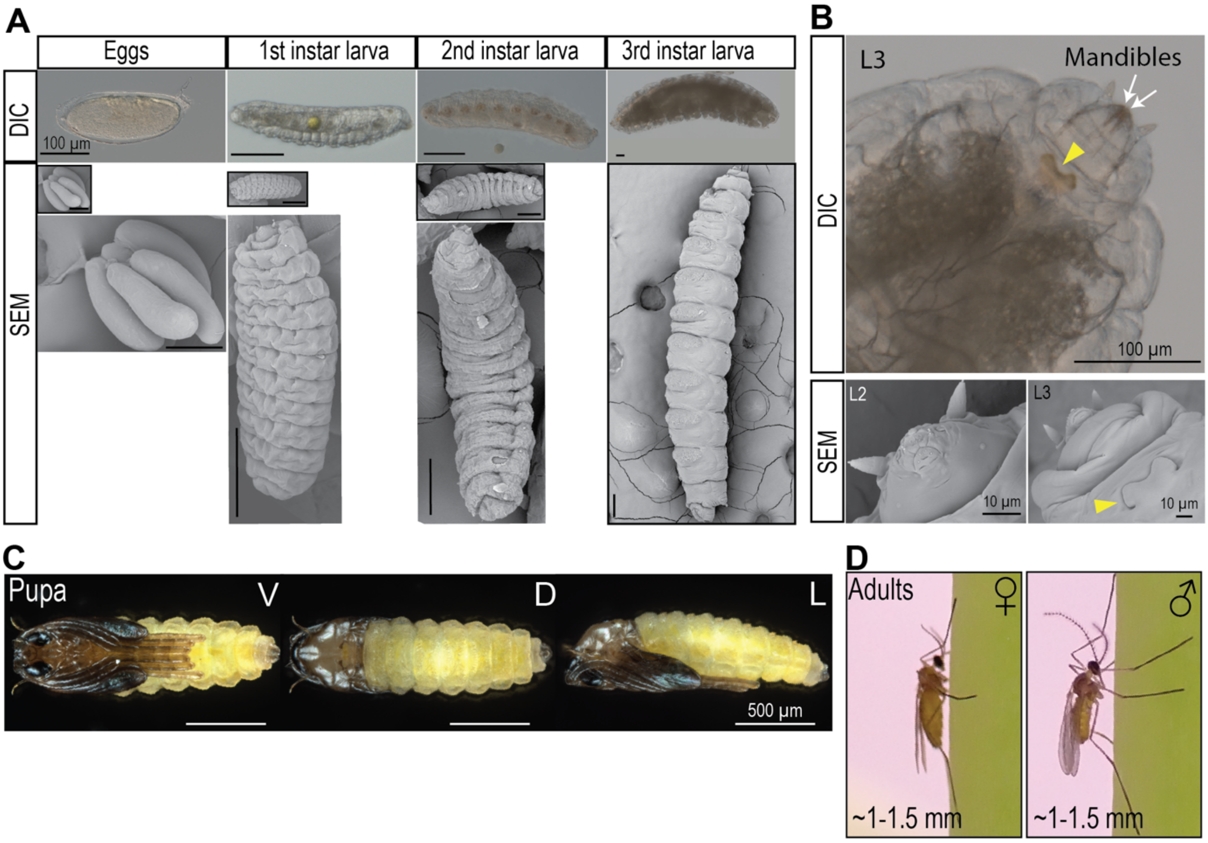


**Fig. S1. Representative images of Swede Midge across life stages.**
**(A)** Differential interference contrast (DIC) and scanning electron microscopy (SEM) images of the egg and the three larval instars (L1, L2 and L3). Top row shows DIC images; bottom row shows SEM images. Insets show enlarged views of eggs, L1 and L2. Scale bars = 100 µm. **(B)** Mouthparts of L2 and L3 larvae. Top: DIC image of L3 larval head. Bottom left: SEM image of L2 mouthparts. Bottom right: SEM image of L3 mouthparts. White arrows indicate mandibles inside the mouth, and yellow arrowheads indicate the spatula. Scale bars: DIC 100 µm; SEM, 10 µm. **(C)** Stereomicroscope images of a Swede Midge pupa in ventral (V), dorsal (D) and lateral (L) views. Scale bars = 500 µm. **(D)** Adult Swede Midges on host plant. Left, female; right, male. Approximate body length excluding wings, antennae and legs is ~1-1.5 mm.

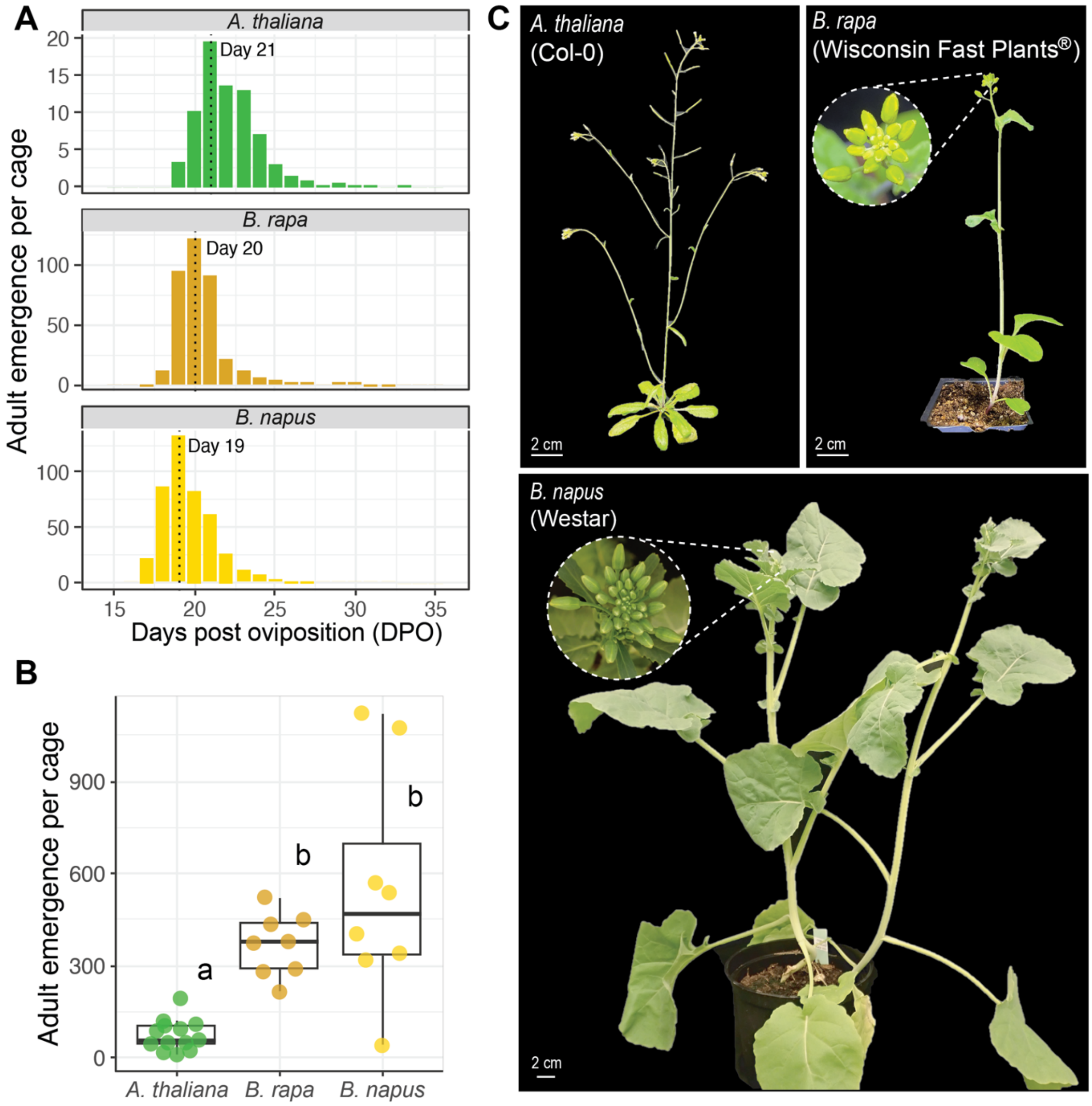


**Fig. S2. Adult SM emergence across Brassicaceae host species.**
Whole-cage infestations were performed using 50 male and 50 female adults, with two plants per cage. **(A)** Adult emergence timing tracking. Mean daily adult emergence per cage for *A. thaliana* Col-0 (13 cages), Rapid Cycling *B. rapa* Wisconsin Fast Plants® (8 cages), and *B. napus* Westar (10 cages), monitored from 15–40 days post oviposition (DPO). Dashed lines indicate peak adult emergence at 21, 20 and 19 DPO, respectively. **(B)** Total adult emergence per cage, summed across all monitored days in (A). Points represent individual cages; boxplots show the median and interquartile range. Different letters indicate significant differences by one-way ANOVA followed by Tukey's HSD test (Tukey-adjusted $P < 0.05$). **(C)** Representative host developmental stages before infestation. *Arabidopsis thaliana* (Col-0) at stages 6.10–6.20 (Boyes *et al.*, 2001); Rapid Cycling *B. rapa* (Wisconsin Fast Plants®) with a young inflorescence and visible closed floral buds with zoomed inset showing inflorescence; and *B. napus* (Westar) at BBCH 57 with zoomed inset showing inflorescence (LANCASHIRE *et al.*, 1991). Scale bars = 2 cm.

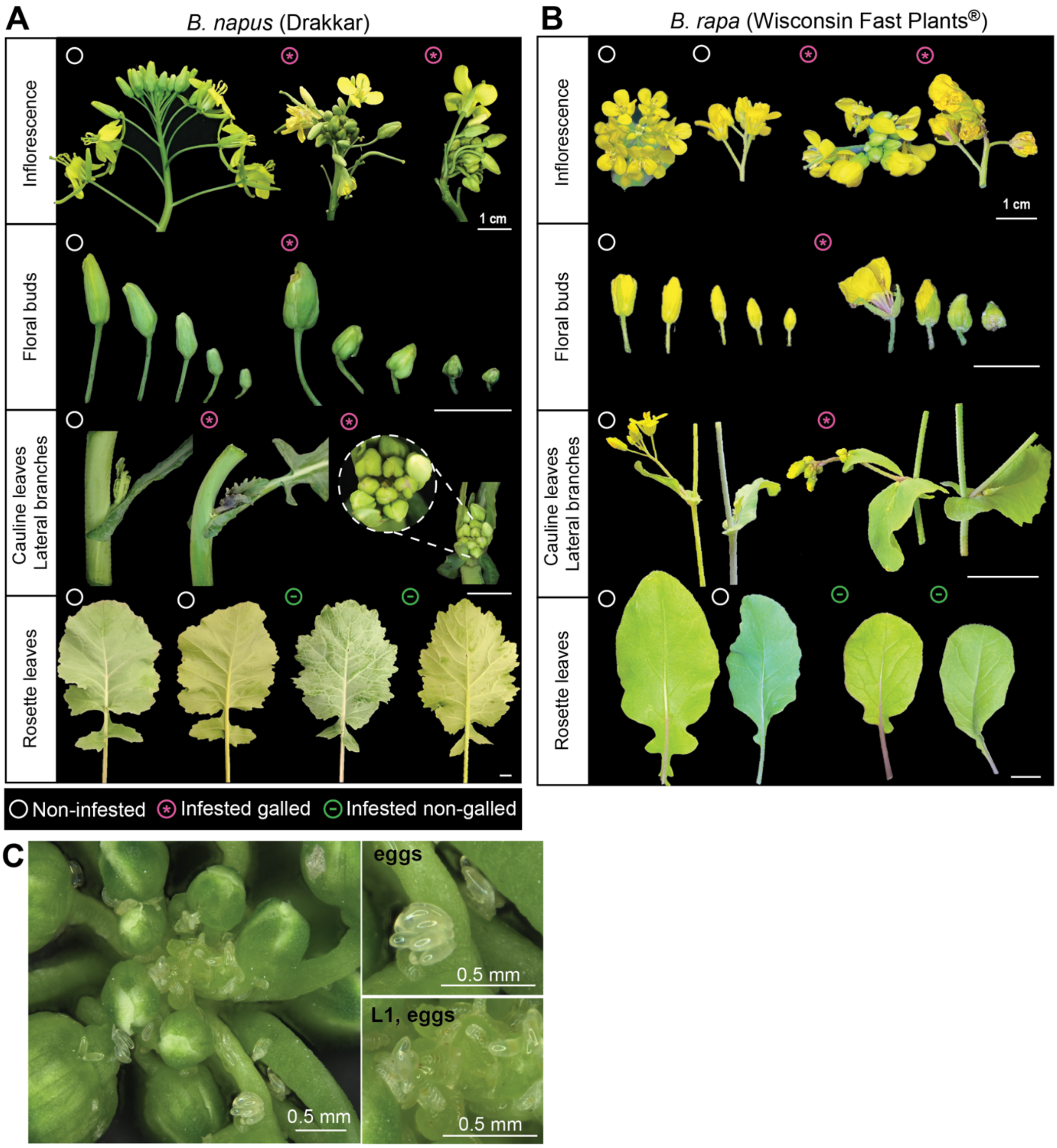


**Fig. S3. Representative Swede Midge gall symptoms on *Brassica* host species.**
**(A)** Representative symptoms on *B. napus* (Drakkar) and **(B)** Rapid Cycling *B. rapa* (Wisconsin Fast Plants®) following whole-plant exposure to Swede Midge adults. Inflorescences, floral buds, cauline leaves and lateral branches and rosette leaves are shown for non-infested controls, infested galled tissues and infested non-galled tissues, as indicated by the symbols. Scale bars = 1 cm. **(C)** Representative images of Swede Midge eggs and L1 larvae on *B. napus* inflorescence and floral tissues following adult infestation. Insets show eggs (top right) and L1 larvae with eggs (bottom right). Scale bars = 0.5 mm.

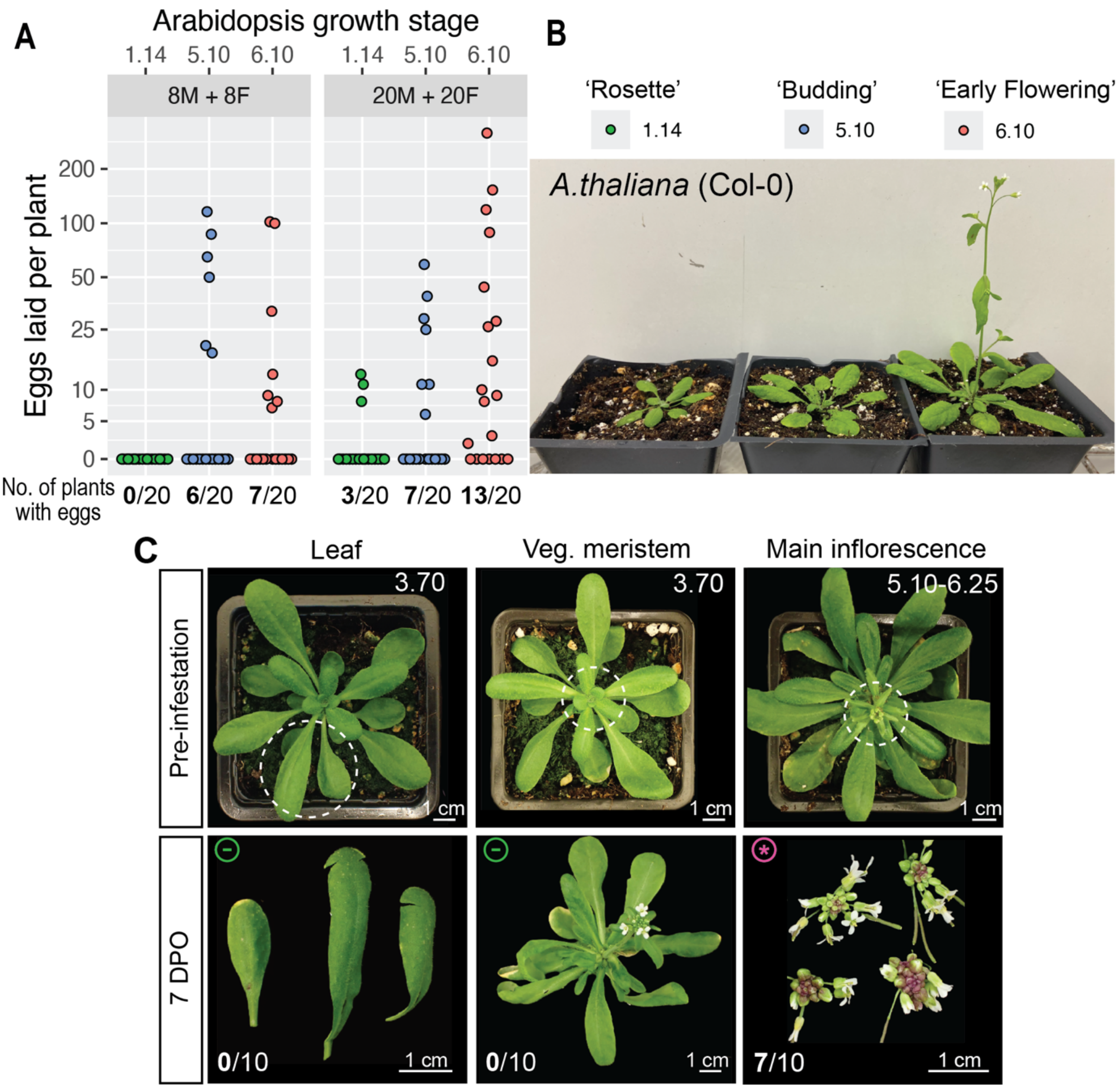


**Fig. S4. Arabidopsis is susceptible to SM at early flowering stages.**

**(A)** Eggs laid per plant on Arabidopsis (Col-0) at different growth stages following whole-cage infestation with either 8M+8F or 20M+20F adults per cage (one plant per cage), with oviposition allowed for 48 h. Points represent egg counts from individual plants; the y-axis is shown on a pseudo-log scale. Numbers for each growth stage indicate plants with eggs/total plants assessed. **(B)** Representative Col-0 plants at growth stages 1.14 (rosette; 14 rosette leaves >1 mm in length), 5.10 (budding; first flower buds visible) and 6.10 (early flowering; ~10% of flowers open) (Boyes *et al.*, 2001) in (A). **(C)** Galling capacity of vegetative tissues. Arabidopsis (Ler-0) rosette leaves or the vegetative meristem at stage 3.70 were infested using clip-cage assays with 5M+5F adults. Main-stem inflorescences served as positive controls using plants from budding (5.10) through early (6.20) and mid-flowering (6.25) stages (Boyes *et al.*, 2001). Representative phenotypes before infestation and at 7 days post oviposition (DPO) are shown. Dashed circles indicate the position at which the clip cage was applied. Numbers for each tissue type indicate plants infested and galled at 7 DPO/total plants assessed. Scale bars = 1 cm.

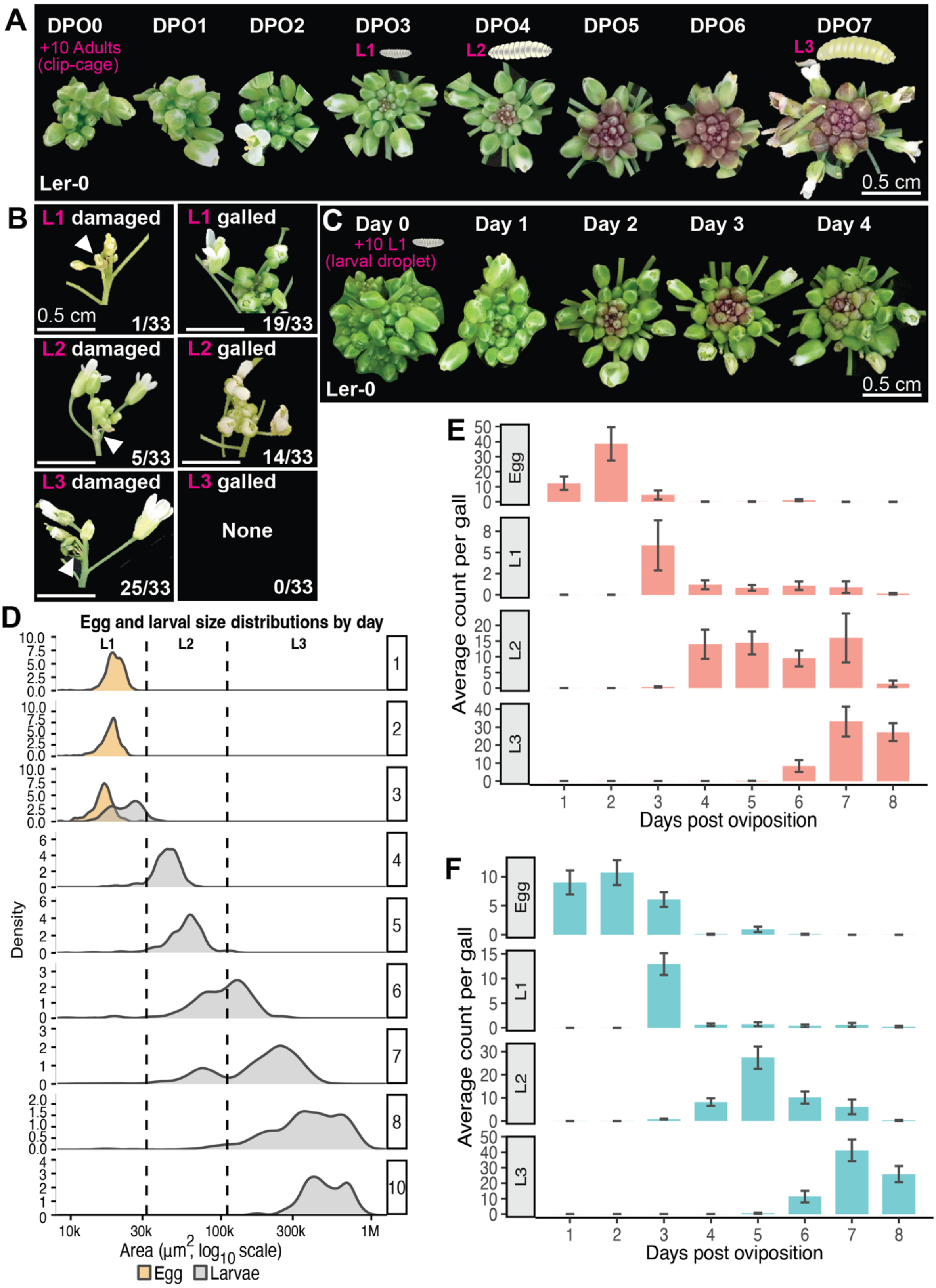


**Fig. S5. Gall and larval development over time following clip-cage and larval droplet infestations.**

**(A)** Daily progression of gall phenotypes from 0–7 days post oviposition (DPO) following clip-cage infestation with 5M+5F adults on Ler-0. This experiment was repeated three times with similar phenotypes. Scale bar = 0.5 cm. **(B)** Representative damaged and galled phenotypes 4 days after application of 5 L1, L2 or L3 larvae to Col-0. Numbers indicate inflorescences displaying the indicated phenotype/total inflorescences assessed (*n* = 33). Arrow heads indicate damaged flower buds. Scale bar = 0.5 cm. **(C)** Daily progression of gall phenotypes of Ler-0 following application of 10 L1 larvae, shown from day 0–4 after larval application. This experiment was repeated three times with similar phenotypes. Scale bar = 0.5 cm. **(D)** Egg and larval area distributions by day and L1–L3 stage classification cutoffs. Density distributions of measured egg and larval area from 1–8 and 10 DPO after clip-cage infestation in Arabidopsis. Data from Col-0 and Ler-0 were combined (*n* = 4,290 segmented objects). Eggs (beige) are shown for days 1–3 as a reference for early L1 developmental size, and larvae are shown in grey. Dashed lines indicate area cutoffs used to classify larval developmental stages: L1, <32,000 $\mu m^2$; L2, 32,000–110,000 $\mu m^2$; and L3, >110,000 $\mu m^2$. **(E,F)** Daily larval developmental stage abundance on Col-0 **(E)** and Ler-0 **(F)** following clip-cage infestation. Using the same dataset summarized in Fig. 3D, bars represent mean numbers of eggs, L1, L2 and L3 per positive sample ± SE at each day post oviposition. Note this is a destructive process and so numbers are not the same between stages. Data were obtained from 94/170 Col-0 and 97/103 Ler-0 infested plants with positive image counts, with 10–15 positive plant replicates per day for each genotype (see also **Table S1**). Larval stages were classified by area cutoffs as described in (D).

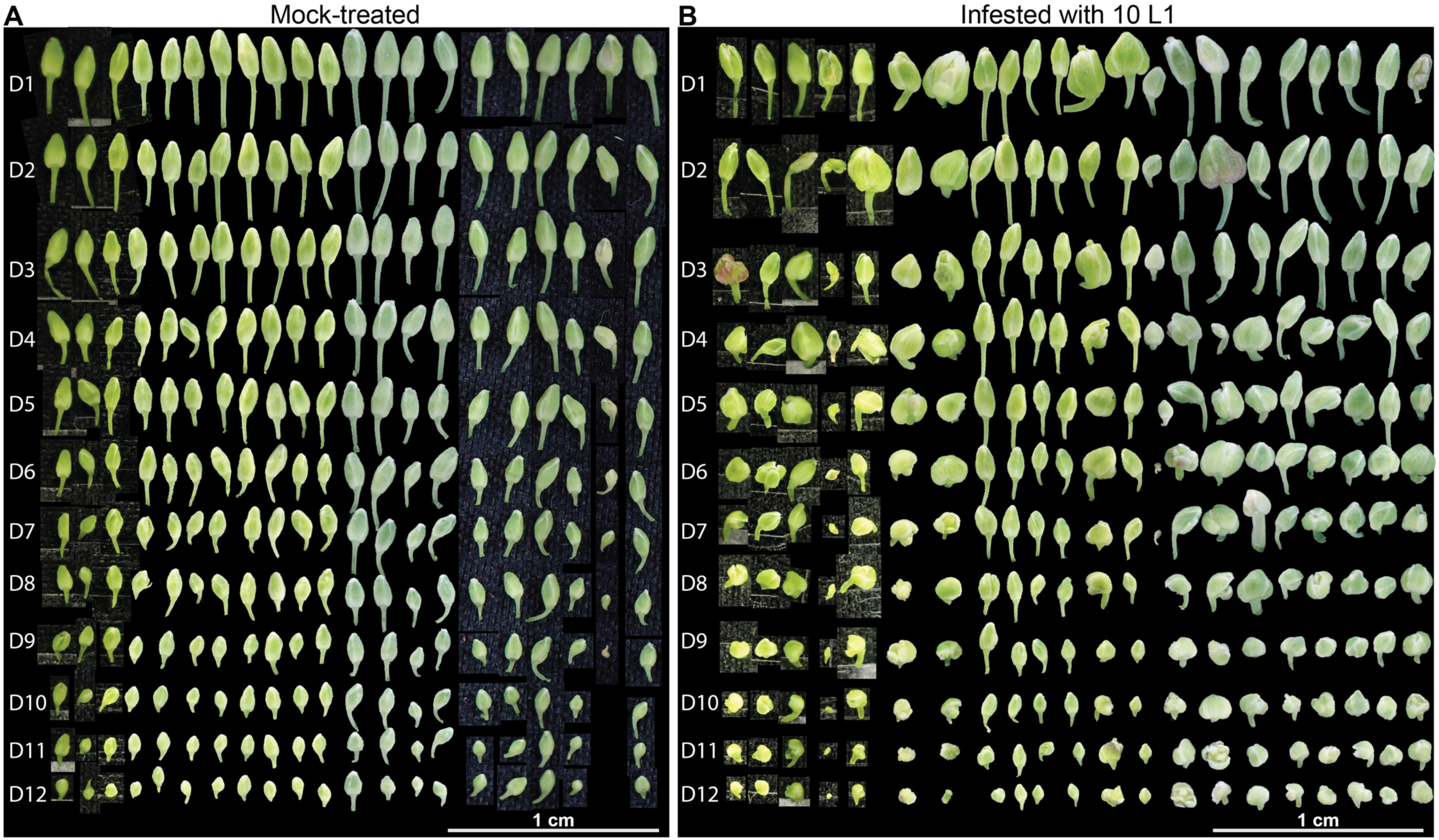


**Fig. S6. Fresh flower buds sampled by phyllotactic dissection.**

Col-0 inflorescences were mock-treated or infested with 10 L1 using larval droplet assays and dissected 4 days after application. Freshly dissected mock-treated **(A)** and infested **(B)** buds are shown. Rows represent dissection orders from D1 (top) to D12 (bottom); each column contains buds dissected from the same inflorescence. Scale bars = 1 cm.

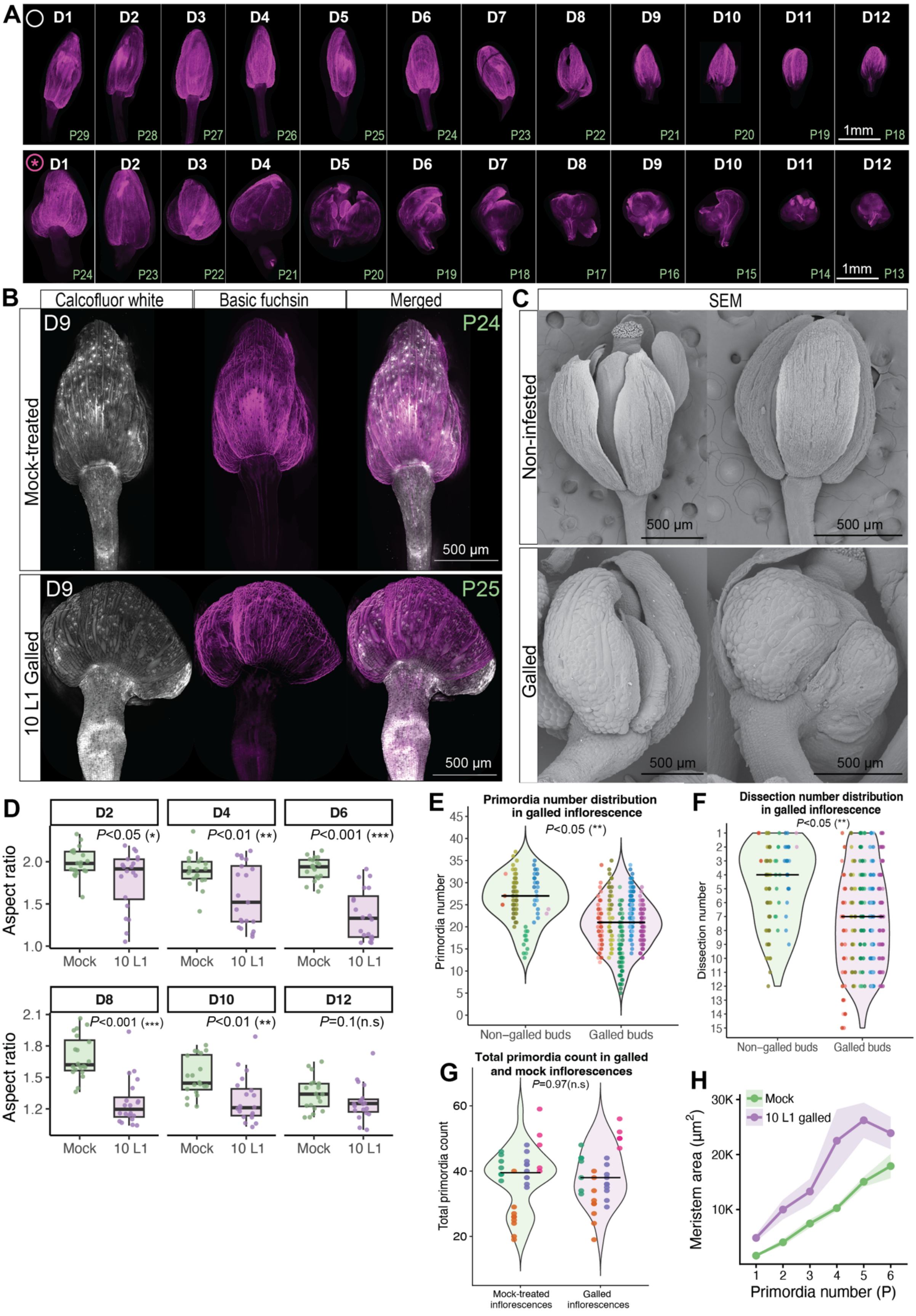


**Fig. S7 Gall-associated changes in floral bud development.**

**(A,B)** Spinning-disk microscopy images of mock-treated and galled floral buds, displayed as maximum-intensity z-projections. Col-0 inflorescences were mock-treated or infested with 10 L1 larvae using larval droplet assays, dissected 4 days after application, and then imaged. White circles indicate mock-treated samples; pink stars indicate infested and galled samples. **(A)** The same D1–D12 buds shown in Fig. 4D; magenta indicates Basic Fuchsin-stained lignified cells. Scale bars = 1 mm. **(B)** Representative D9 buds; Calcofluor White-stained cell walls are shown in grey and Basic Fuchsin staining in magenta. D and P labels indicate dissection order and primordium number, respectively. Scale bars = 500 µm. **(C)** Representative SEM images of non-infested and galled floral buds. Scale bars = 500 µm. **(D)** Bud aspect ratio was quantified from photographs of freshly dissected buds from mock-treated and 10 L1 treated samples at D2, D4, D6, D8, D10, and D12 (see raw images in Fig. S6). Each dissection order included 20–21 mock-treated plants and 19–21 10 L1 treated plants. Points represent individual buds; boxplots show the median and interquartile range. *P*-values are from Welch's *t*-tests comparing mock-treated and 10 L1 treated groups within each dissection. **(E,F)** Violin plots showing the distributions of primordium number **(E)** and dissection order **(F)** for non-galled ($n$ = 85) and galled ($n$ = 282) buds from 36 galled inflorescences across five experiments. Galled inflorescences were defined as those containing at least one galled bud among D1–D12. Points represent individual buds, colours indicate individual plants grouped by experiment and black horizontal bars indicate medians. Paired plant-level medians were compared using two-sided Wilcoxon signed-rank tests ($P$ = 0.0059 for both E and F). **(G)** Violin plot showing the distributions of total primordia counts in galled ($n$ = 33) and mock-treated ($n$ = 32) inflorescences. All open flowers were cut prior to treatment. Total primordia count after infestation represents the sum of siliques, open flowers and floral buds counted to the apex of each inflorescence 4 days after application. Points represent individual inflorescences; colours indicate individual plants grouped by experiment. Black horizontal bars indicate medians. Total primordia counts were compared using a two-sided Wilcoxon rank-sum test ($P$ = 0.97). **(H)** Mean bud area across primordia series (P1-P6), shaded area indicates ± SE. Sample sizes: Mock $n$ = 121, 10 L1 galled $n$ = 109.

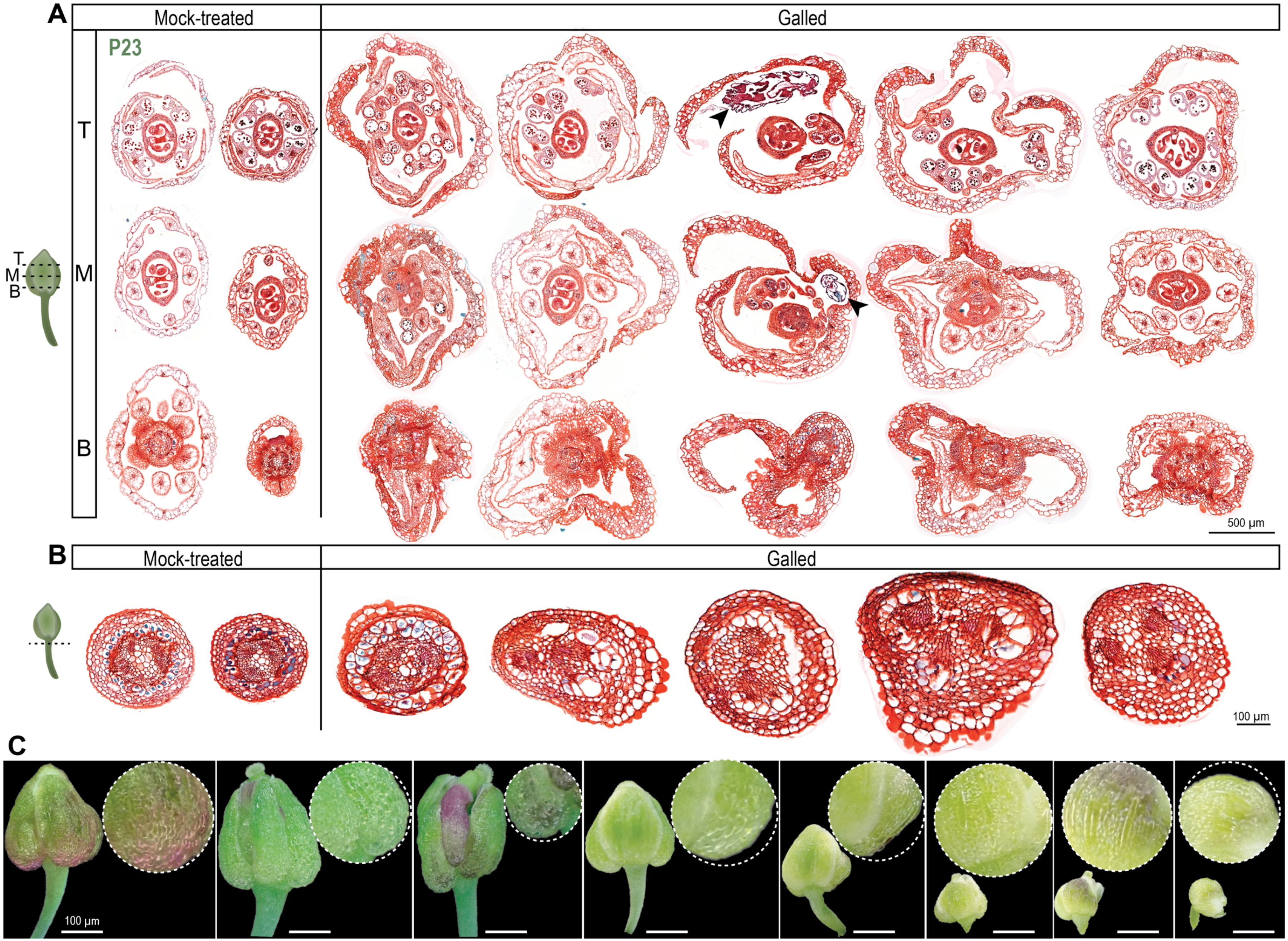

A
Mock-treated
Galled
P23
T
M
B
500 µm
B
Mock-treated
Galled
100 µm
C
100 µm

**Fig. S8. Additional gall-associated morphological and histological phenotypes.** Col-0 inflorescences were mock-treated or infested with 10 L1 larvae using larval droplet assays and analysed 4 days after application. **(A)** Additional replicates of the floral bud histology shown in Fig. 6C. Transverse sections from mock-treated and galled P23 floral buds were stained with Safranin O and Fast Green. Sections from the same bud are arranged vertically and correspond to the top (T), middle (M), and base (B) positions indicated in the bud schematic. Arrowheads indicate Swede Midge larvae. Scale bar = 500 µm. **(B)** Additional replicates of the pedicel histology shown in Fig. 6D. Transverse sections from mock-treated and galled pedicels were stained with Safranin O and Fast Green. Scale bar = 100 µm. **(C)** Representative fresh galled floral buds arranged from older to younger developmental positions from left to right, showing gall-associated "bubbly" cells visible at the sepal surface. Dashed circles show magnified views of the corresponding sepal regions. Scale bars = 100 µm.

### Supplemental Table Legends

**Table S1. Sample-level SM egg and larval populations during development on Col-0 and Ler-0.** Individual clip-cage-infested inflorescences were sampled from 1–8 DPO (*n* = 273). Each row represents one inflorescence and reports ecotype, DPO, gall status, numbers of eggs and L1–L3 larvae recovered, total larval number, mean larval area and summed larval area. These samples underlie the analyses in **Fig. 3D**, **Fig. S5E–F.**

**Table S2. Egg recovery from Col-0 and Ler-0 following adult SM infestation.** Clip-cage infested plants were sampled. For each ecotype and DPO, the table reports the total number of infested plants sampled, the number and percentage of plants from which at least one egg was recovered, the total number of eggs recovered and the mean egg number per egg-positive plant. Mean egg number was calculated only among egg-positive plants. Values were summarised from the same developmental dataset used for Table S1.
